# Bridging the gap between presynaptic hair cell function and neural sound encoding

**DOI:** 10.1101/2022.10.19.512823

**Authors:** Lina María Jaime Tobón, Tobias Moser

## Abstract

Neural diversity can expand the encoding capacity of a circuitry. A striking example of diverse structure and function is presented by the afferent synapses between inner hair cells (IHCs) and spiral ganglion neurons (SGNs) in the cochlea. Presynaptic active zones at the pillar IHC side activate at lower IHC potentials than those of the modiolar side that have more presynaptic Ca^2+^-channels. The postsynaptic SGNs differ in their spontaneous firing rates, sound thresholds and operating ranges. While a causal relationship between synaptic heterogeneity and neural response diversity seems likely, experimental evidence linking synaptic and SGN physiology has remained difficult to obtain. Here, we aimed at bridging this gap by *ex vivo* paired recordings of IHCs and postsynaptic SGN boutons with stimuli and conditions aimed to mimic those of *in vivo* SGN-characterization. Synapses with high spontaneous rate of release (*SR*) were found predominantly on the pillar side of the IHC. These high *SR* synapses had larger and more temporally compact spontaneous EPSCs, lower voltage-thresholds, tighter coupling of Ca^2+^ channels and vesicular release sites, shorter response latencies and higher initial release rates. This study indicates that synaptic heterogeneity in IHCs directly contributes to the diversity of spontaneous and sound-evoked firing of SGNs.

**Significance Statement:** Sound encoding relies on spiral ganglion neurons (SGNs) with diverse spontaneous firing, sound thresholds of firing and sound-intensity range over which SGN firing rate changes. Such functional SGN diversity might originate from different input from afferent synapses with inner hair cells (IHCs). The present study addresses this hypothesis by using recordings from individual IHC-SGN synapses of hearing mice under *ex vivo* conditions aimed to mimic cochlear physiology. The results provide evidence that synaptic heterogeneity in IHCs contributes to SGN firing diversity. Thus, the cochlea employs heterogeneous synapses to decompose sound information into different neural pathways that collectively inform the brain about sound intensity.

## Introduction

Chemical synapses represent diverse and plastic neural contacts that are adapted to the specific needs of neural computation. Synaptic diversity is expressed across the nervous system, within a given circuit and even within the same neuron (recent reviews in Ref. Nusser, 2018; Wichmann and Kuner, 2022). Synaptic diversity occurs at various levels: from synapse shape and size, to the ultrastructure of pre- and postsynaptic specializations, to their molecular composition. The auditory system harbors striking examples of synaptic diversity. Glutamatergic ribbon synapses in the cochlea, calyceal synapses in the brainstem, and bouton synapses throughout the central auditory system differ greatly from each other (Moser et al., 2006; Wichmann and Kuner, 2022). Beyond the diversity across synapses formed by different neurons and at different places (e.g. different regions of the brain or frequency (tonotopic) places of the cochlea (Johnson et al., 2017)), heterogeneity exists even among auditory synapses formed by an individual presynaptic inner hair cell (IHC) with its 5-30 postsynaptic spiral ganglion neurons (SGNs, (reviews in: Gómez-Casati and Goutman, 2021; Meyer and Moser, 2010; Moser, 2020; Reijntjes and Pyott, 2016)). Synaptic heterogeneity has been found at different tonotopic positions of the cochlea and is a candidate mechanism for how the cochlea decomposes acoustic information (Grant et al., 2010; Meyer et al., 2009; Ohn et al., 2016; Özçete and Moser, 2021). For example, the cochlea might use heterogeneous afferent synapses to break down sound intensity information into complementary spike rate codes of SGNs that have been reported along the tonotopic axis of the cochlea for several species (Huet et al., 2016; Kiang et al., 1965; Sachs and Abbas, 1974; Taberner and Liberman, 2005; Winter et al., 1990).

Decades of *in vivo* recordings from single SGNs have demonstrated functional diversity of SGNs with comparable frequency tuning, i.e., receiving input from IHCs at a given tonotopic place or potentially even the same IHC. Such functional diversity is present in both spontaneous and sound-evoked firing. The spontaneous firing rate (SR) in the absence of sound varies from less than 1 spikes/s to more than 100 spikes/s (Barbary, 1991; Evans, 1972; Kiang et al., 1965; Schmiedt, 1989; Taberner and Liberman, 2005). In response to increasing sound pressure, SGNs with high SR show a low sound threshold and a steep rise in the spike rate to increasing sound intensities until the rate saturates. SGNs with low SR have a higher sound thresholds, shallower spike rate rise and saturate at higher sound intensities (Ohlemiller et al., 1991; Winter et al., 1990). Additionally, SGNs show differences in their excitability (Crozier and Davis, 2014; Markowitz and Kalluri, 2020; Smith et al., 2015), morphological features (Liberman, 1980; Merchan-Perez and Liberman, 1996; Tsuji and Liberman, 1997) and heterogeneous molecular profiles (Li et al., 2020; Petitpré et al., 2020, 2018; Shrestha et al., 2018; Sun et al., 2018). Yet, it has remained challenging to demonstrate a causal link of a candidate mechanism to the physiological SGN diversity.

One common approach has been to capitalize on a pioneering study that employed *in vivo* labelling of physiologically characterized SGNs in cats and proposed that synapses formed by low and high SR SGNs segregate on the basal IHC pole (Liberman, 1982). High SR SGNs preferentially contacted the pillar side of the IHC (facing pillar cells), while low SR SGNs synapsed on the opposite, modiolar side of the IHC (facing the cochlear modiolus). Interestingly, a segregation has also been found for afferent and efferent synaptic properties, as well as molecular and biophysical SGN properties (Frank et al., 2009; Grant et al., 2010; Hua et al., 2021; Kantardzhieva et al., 2013; Liberman et al., 2011; Markowitz and Kalluri, 2020; Merchan-Perez and Liberman, 1996; Meyer et al., 2009; Neef et al., 2018; Ohn et al., 2016; Özçete and Moser, 2021; Shrestha et al., 2018; Sun et al., 2018; Yin et al., 2014). For instance, type I_b_ and I_c_ SGNs preferentially synapse on the modiolar side (Sherrill et al., 2019; Shrestha et al., 2018; Sun et al., 2018) and show low SR (Siebald et al., 2023). Pillar synapses are preferentially formed by type I_a_ SGNs (Shrestha et al., 2018; Siebald et al., 2023), have smaller IHC active zones (AZs) (Liberman et al., 2011; Ohn et al., 2016; Reijntjes et al., 2020), and activate at voltages as low as the IHC resting potential (Ohn et al., 2016; Özçete and Moser, 2021). The low voltage of activation of release at pillar synapses shown *ex vivo* could underly the high SR and low sound threshold of firing of SGNs found *in vivo* but direct demonstration of such a link has been missing.

Here, we aimed to bridge the gap between *ex vivo* presynaptic physiology and *in vivo* SGN neurophysiology. We performed paired IHC and SGN bouton recordings in acutely explanted organs of Corti from hearing mice with stimuli and conditions aimed to mimic *in vivo* SGN-characterization. This approach tightly controls IHC Ca^2+^ influx and records the postsynaptic SGN response to glutamate release at a single afferent synapse. Using the rate of spontaneous excitatory postsynaptic currents (rate of sEPSCs, *SR*) as a surrogate for SGN SR, we demonstrate that high *SR* synapses have larger and more compact sEPSCs as well as lower voltage thresholds, shorter latencies of evoked EPSCs, tighter Ca^2+^ channel coupling to vesicle release and higher initial release rates. 90% of these high *SR* synapses were located on the pillar side of the IHC. Our findings suggest that synaptic heterogeneity accounts for much of the SGN firing diversity at a given tonotopic position.

## Results

Simultaneous paired patch-clamp recordings from IHCs and one of the postsynaptic SGN boutons were performed on mice after the onset of hearing (postnatal days (p) 14-20). We performed perforated-patch whole-cell configuration from IHCs, held at their presumed physiological resting potential (−58 mV (Johnson, 2015)), and ruptured-patch whole-cell recordings from one of the postsynaptic SGN boutons (Fig. 1). Due to the technical difficulty of establishing the paired recording, typically only one bouton was recorded per IHC. Recordings were made at body temperature and in artificial perilymph-like solution (Wangemann and Schacht, 1996). To establish the paired recording, we approached boutons facing either the pillar or the modiolar side of the IHC in an effort to elucidate synaptic differences between both sides (Fig. 1A). We nickname the synapse location as “pillar” and “modiolar” based on the DIC-image, but note that efforts to stain and image the recorded boutons by fluorescence microscopy were not routinely successful. In addition, the recorded boutons were classified based on their spontaneous rate of synaptic transmission (Fig. 1B, Fig. 2 and related figure supplements; *SR,* **Low *SR*** < 1 sEPSC/s vs **High *SR*** ≥ 1 sEPSC/s according to (Taberner and Liberman, 2005)). We then performed an *in-depth* biophysical analysis of evoked release (Fig. 1C and Figs. 3 – 4, and related figure supplements).

**Figure 1.**
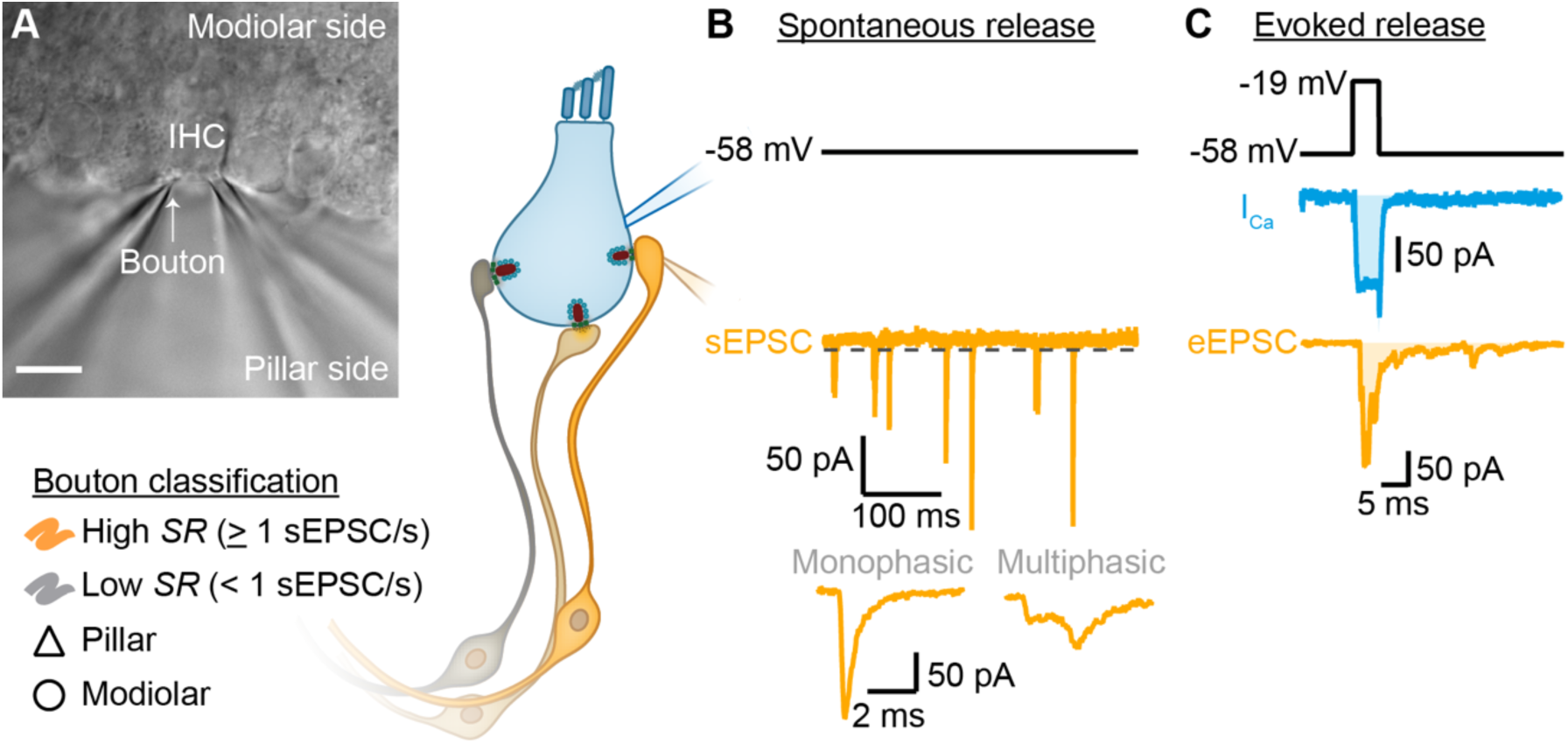
Paired IHC-bouton patch-clamp recordings to study the release properties of individual IHC ribbon synapses as a function of synapse position. **A.** Differential Interference Contrast image of an explanted murine organ of Corti. In this example, supporting cells from the pillar side were removed to gain access to the IHCs and their contacting boutons. The recorded boutons were classified based on their position (△ pillar or ❍ modiolar) and on their spontaneous rate (*SR*) (Low *SR* < 1 sEPSC/s vs High *SR* ≥ 1 sEPSC/s). Scale bar: 10 µm. **B.** Spontaneous release was recorded in absence of stimulation (i.e., IHC holding potential = −58 mV; Supplementary Table 1; dashed line represents the threshold for sEPSC detection). sEPSCs were classified as monophasic (a steady rise to peak and monoexponential decay, temporally more compact) or as multiphasic (multiple inflections and slowed raising and decaying kinetics, non-compact). **C.** Evoked release: depolarizing pulses (black trace) were used to trigger whole IHC Ca^2+^ influx (I_Ca_, blue trace) and ensuing release of neurotransmitter that evoked EPSCs (eEPSCs, light orange trace). Ca^2+^ charge and eEPSC charge were estimated by taking the integral of the currents (shaded light blue and light orange areas).

**Figure 2.**
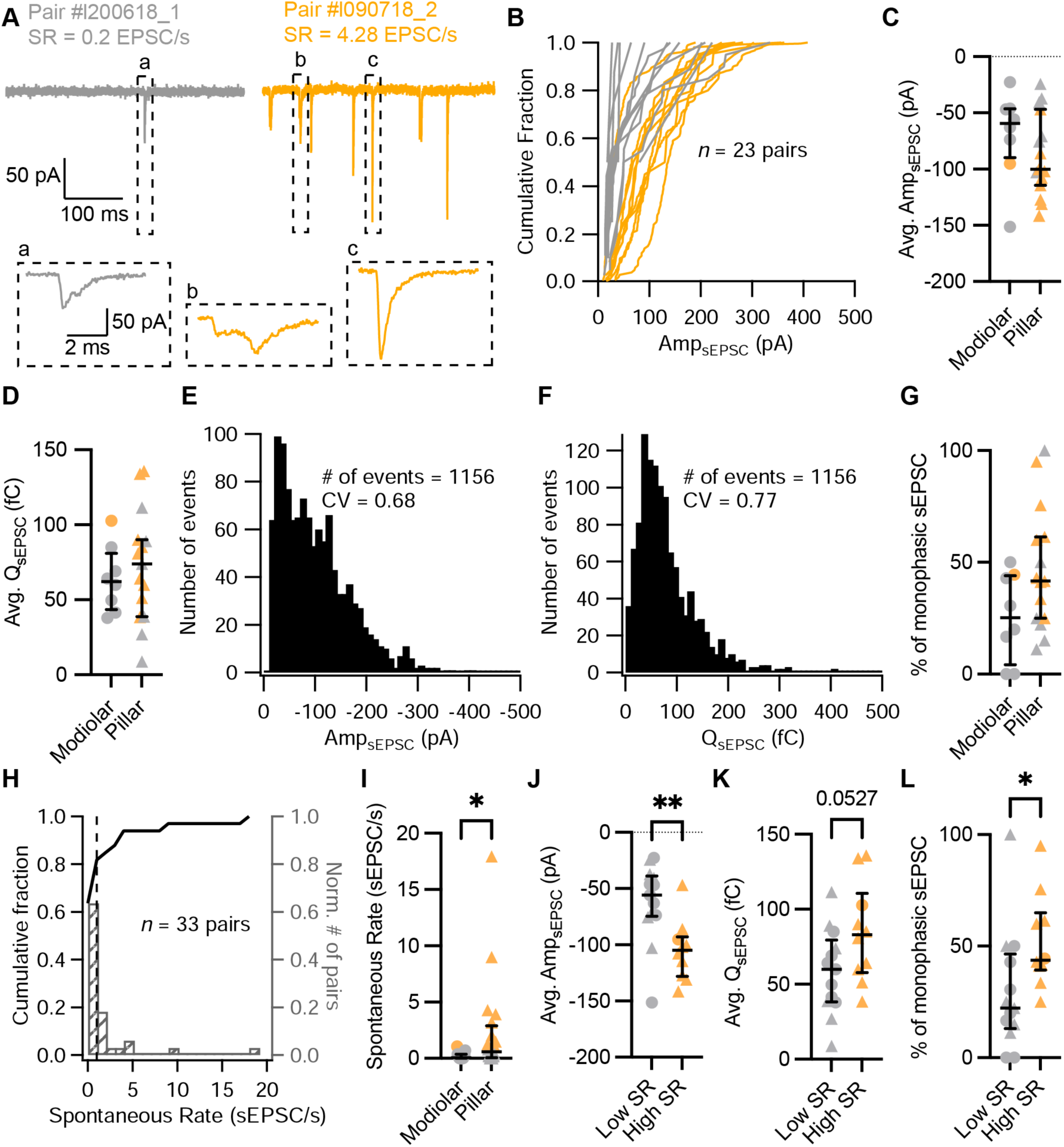
Synapses with high spontaneous release have larger and more monophasic sEPSCs. **A.** Spontaneous EPSCs recorded in the absence of stimulation (i.e., IHC holding potential = −58 mV) from two exemplary paired recordings with different spontaneous rate (*SR:* grey for low *SR,* orange for high *SR*). ‘Pair #’ identifies individual paired recordings. Insets show the selected sEPSCs in an expanded time scale. (a,b) correspond to multiphasic sEPSCs, while (c) represents a typical monophasic sEPSC. **B.** Cumulative sEPSC amplitude plots for 23 paired synapses that had spontaneous release. **C-D.** Average sEPSC amplitude (C) and charge (D) from individual synapses recorded from the pillar or modiolar side of the IHC. **E-F.** Pooled sEPSC amplitude (E) and charge (F) distributions show a distinct peak at −40 pA and 40 pC, respectively. Bin size: 10 pA or pC. **G.** Percentage of monophasic sEPSCs in pillar and modiolar synapses. **H.** Cumulative fraction (left axis) and normalized histogram (right axis) of the spontaneous rate (bin size is 1 sEPSC/s) of 33 pairs. **I.** Pillar synapses had higher rates of sEPSCs. **J-L.** High *SR* synapses had significantly larger sEPSC amplitudes (J), a tendency to bigger sEPSC charges (K) and higher percentages of monophasic sEPSCs (L). Panels G, I-L show individual data points with the median and interquartile range overlaid (line). Synapses were classified as △ pillar or ❍ modiolar, and as Low *SR* < 1 sEPSC/s ≤ High *SR*.

**Figure 3.**
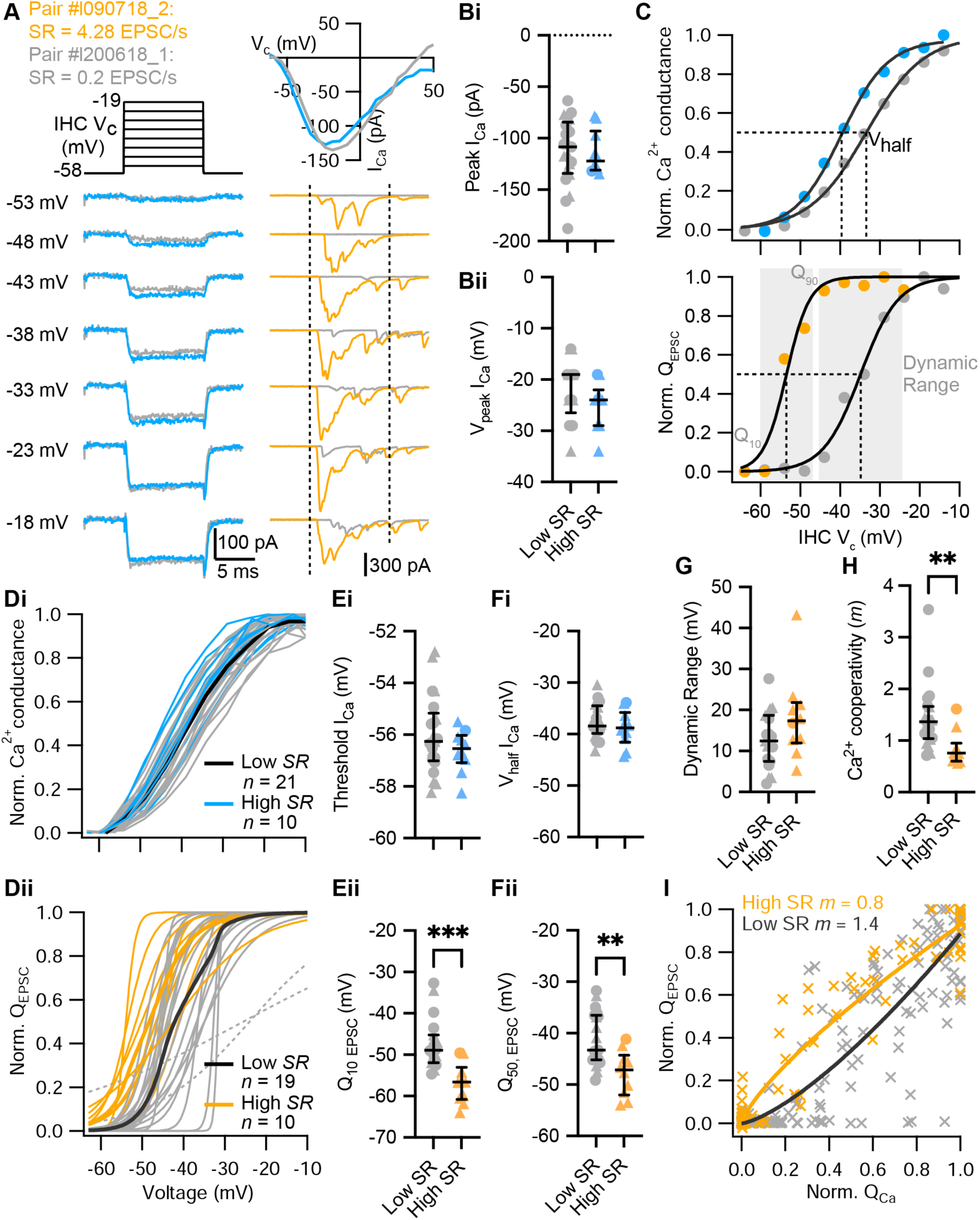
High *SR* synapses activate at lower voltages and show tighter Ca^2+^ channel coupling of synaptic release. **A.** Voltage-protocol (top left), IHC Ca^2+^ current (I_Ca_, bottom left and top right, blue and gray) and eEPSCs (bottom right, orange and gray) of a high and a low *SR* pair, respectively, in response to 10 ms depolarizations (dashed vertical lines on top of eEPSC data indicate the onset and offset of the depolarization) to different potentials ranging from −58 to −19 mV in 5 mV steps. The upper right panel shows the current-voltage relationships for the two pairs. **Bi-Bii.** The peak of whole-cell Ca^2+^ current (Bi) and the voltage eliciting maximum Ca^2+^ current (Bii) of IHCs were comparable between high and low *SR* synapses. **C.** Upper panel: Fractional activation of the Ca^2+^ channels (blue and gray data points from the examples shown in A) was obtained from the normalized chord conductance. Voltage of half-maximal activation (V_half Ca_; dotted line) and voltage sensitivity of activation (slope *k*) were determined using a Boltzmann fit (black trace) to the activation curve. Lower panel: Release-intensity curve (orange and gray data points from the examples shown in A) was obtained from the Q_EPSC_ for each depolarization step. A sigmoidal function (black trace) was fitted to obtain the voltage of half-maximal synaptic release (V_half_ Q_EPSC_; dotted line) and the voltage sensitivity of the release (slope), as well as the dynamic range for which the exocytosis changes from 10-90% (gray area). **Di-Dii.** Voltage dependence of whole-cell Ca^2+^ channel activation (activation curve; Di) and fits to release-intensity curves (Dii) for 31 synapses. Averages (thick lines) and individual curves (thin lines) are overlaid. The release-intensity curve of two low *SR* pairs could not be fitted (grey dotted lines). **Ei-Fi.** The threshold of Ca^2+^ influx (Ei) and V_half_ I_Ca_ (Fi) did not differ between low and high *SR* synapses. **Eii-Fii**. Voltage of 10% maximum release (Q_10 EPSC_, Eii) and V_half_ Q_EPSC_ (Fii) were significantly more hyperpolarized in high *SR* synapses. **G.** Dynamic range of release was comparable between low and high *SR* synapses. **H.** Ca^2+^ cooperativity (*m*) estimated from fitting a power function to the Q_EPSC_ – Q_Ca_ relationship for each individual synapse (see Figure 3–figure supplement 2) was significantly lower in high *SR* synapses. **I.** Scatter plot of normalized Q_EPSC_ versus the corresponding normalized Q_Ca_. The solid lines are a least-squares fit of a power function (Q_EPSC_ = a(Q_Ca_)*^m^*) to the data yielding *m*_high *SR*_ of 0.8 and *m*_low *SR*_ of 1.4. Panels B, E-H show individual data points with the median and interquartile range overlaid (line). Synapses were classified as △ pillar or ❍ modiolar, and as Low *SR* < 1 sEPSC/s ≤ High *SR*.

### Spontaneous synaptic transmission

In order to recapitulate synaptic transmission in the absence of sound stimulation, we held the IHC at their presumed physiological resting potential (−58 mV (Johnson, 2015)). Since Ca_V_1.3 Ca^2+^-channels activate at low voltages (−65 to −45 mV (Koschak et al., 2001; Picher et al., 2017; Platzer et al., 2000; Xu and Lipscombe, 2001)), their open probability at the IHC resting potential is thought to be sufficient to trigger spontaneous release (Glowatzki and Fuchs, 2002; Özçete and Moser, 2021; Robertson and Paki, 2002). Under our experimental conditions, spontaneous, i.e., excitatory postsynaptic currents in the absence of IHC stimulation (sEPSCs) were observed in 23 of 33 pairs (Fig. 1B, Fig. 2, 20 recordings were targeted to the pillar side and 13 to the modiolar side). We used the rate of sEPSCs (*SR)* as a surrogate of SGN SR as sEPSCs trigger SGN firing with >90% probability (Rutherford et al., 2012). Regardless of all efforts to maintain the physiological integrity in the *ex vivo* experiments, we expect our estimated rates of sEPSC to underestimate the SGN SR for the same age group (Wong et al., 2013). For each paired recording, we quantified the *SR* during 5 or 10 s of a continuous recording, or during the segment before and after the step depolarization protocols (Supplementary Table 1). Amplitudes of sEPSCs typically varied from around −10 pA to −400 pA across all recorded IHC-SGN synapses (Fig. 2B). The amplitude histogram for all pairs was slightly skewed towards smaller amplitudes (skewness of 1.06) with a coefficient of variation (CV) of 0.68 (a similar distribution was obtained from 12 bouton-only recordings during which the IHC was not patched [Figure 2–figure supplement 1, panel A]). The charge distribution for all pairs displayed a prominent peak at 40 fC, with a skewness of 2.00 and a CV of 0.77. *SR* ranged from 0 to about 18 sEPSC/s (Fig. 2H; a similar range of 0 to about 16 sEPSC/s was recorded without patch-clamping the IHC). The *SR* distribution was highly skewed, with a median of 0.2 sEPSC/s. Following the spontaneous firing rate classification of mouse SGNs by Taberner and Liberman (2005), we classified the synapses into low (< 1 sEPSC/s, ∼70%) or high (≥ 1 sEPSC/s, ∼30%) *SR* synapses (see Fig. 2A for examples). Next, we analyzed the recordings as a function of i) synapse position and ii) rate of spontaneous synaptic transmission.

### i) Position dependence of synaptic transmission

Of the 33 obtained paired recordings, 20 were classified as pillar synapses. The mean amplitude (Fig. 2C) and charge (Fig. 2D) of the sEPSCs were comparable between modiolar and pillar SGN boutons (Amp_sEPSC_ −69.37 ± 13.91 pA [*n* = 8 modiolar] vs −87.21 ± 9.49 pA [*n* = 15 pillar]; *p* = 0.2913, unpaired t-test; Q_sEPSC_ 63.74 ± 7.77 fC [modiolar] vs 72.54 ± 9.57 fC [pillar]; *p* = 0.5463, unpaired t-test). sEPSC of pillar SGN boutons showed significantly shorter 10-90% rise times than the modiolar ones (Figure 2–figure supplement 1, panel B and C; 0.38 ± 0.04 ms vs 0.57 ± 0.06 ms; *p* = 0.0111, unpaired t-test), yet similar decay times and full-width half-maximum (FWHM, Figure 2– figure supplement 1, panels B, D-E; *p* = 0.7997 and *p* = 0.9198, respectively, unpaired t-test). As a second approach to sEPSC properties, we quantified the percentage of monophasic (or temporally more compact) sEPSCs (Chapochnikov et al., 2014; Glowatzki and Fuchs, 2002) and found a non-significant trend towards higher percentages of monophasic sEPSCs for pillar synapses (Fig. 2G; 25.57 ± 6.9% for modiolar vs 46.65 ± 7.06% for pillar; *p* = 0.0681, unpaired t-test). Finally, *SR* was significantly higher for the pillar synapses (Fig. 2I, mean *SR* of 2.30 ± 0.96 sEPSC/s; median 0.59; *n* = 20) compared to modiolar ones (Fig. 2I, mean *SR* of 0.22 ± 0.09 sEPSC/s; median 0.05; *n* = 13; *p* = 0.0311, Mann-Whitney U test).

### ii) Relation of synaptic properties to the rate of spontaneous synaptic transmission

High *SR* synapses had significantly larger sEPSCs (Fig. 2J and Figure 2–figure supplement 1, panel Fi; average sEPSC amplitude of −105.2 ± 8.47 pA for high *SR* [*n* = 10] vs. −62.39 ± 9.67 pA for low *SR* [*n* = 13]; *p* = 0.0042, unpaired t-test). sEPSC charge tended to be larger in high *SR* SGN synapses (Fig. 2K and Figure 2–figure supplement 1, panel Fii; Q_sEPSC_ 84.23 ± 10.38 fC for high *SR* vs. 58.14 ± 7.79 fC for low *SR*; *p* = 0.0527, unpaired t-test). The fraction of monophasic sEPSCs was significantly higher in high *SR* synapses (Fig. 2L; 52.08 ± 6.65%; median 43.71%) compared to low *SR* synapses (29.49 ± 7.43%; median 22.22%; *p* = 0.0185, Mann-Whitney U test). High *SR* synapses also showed a significantly faster 10-90% rise times (Figure 2–figure supplement 1, panel G; 0.36 ± 0.04 ms) than low *SR* synapses (0.51 ± 0.05 ms; *p* = 0.0420, unpaired t-test).

Other sEPSCs kinetics, such as decay time constant and FWHM, were not different between low and high *SR* pairs (Figure 2–figure supplement 1, panels H, I; *p* = 0.7969 and *p* = 0.9948, respectively, unpaired t-test). Taken together, these results indicate that high *SR* synapses are characterized by sEPSCs with larger amplitudes, faster rising times and a more compact waveform, while significant differences of pillar and modiolar synapses were limited to sEPSC rise times. Yet, 9 out of 10 synapses with *SR* ≥ 1 sEPSC were located on the pillar side of the IHC.

### Evoked synaptic transmission differs between afferent synapses with high and low *SR*

Next, we compared the physiology of afferent synapses with high and low *SR* by adapting stimulation protocols routinely employed for *in vivo* characterization of sound encoding by SGNs. We used step depolarizations to emulate physiological receptor potentials given that mature IHCs of the “high frequency” mouse cochlea have graded receptor potentials that primarily represent the rectified envelope of an acoustic stimulus (i.e. the DC component (Russell and Sellick, 1978)).

### i) Stimulus intensity encoding at IHC synapses

Sound intensity encoding by SGNs primarily relies on a spike rate code: the average discharge rate increases with the strength of the acoustic stimuli from threshold to saturation of the response. These so-called “rate level functions” are typically analyzed by fitting a sigmoidal function, of which the range of sound pressure level between 10 and 90% of the maximal discharge rate represents the operational or dynamic range (Sachs and Abbas, 1974; Taberner and Liberman, 2005; Winter et al., 1990). To understand stimulus intensity coding at mouse IHC synapses, we measured whole-cell IHC Ca^2+^ currents and the evoked EPSCs (eEPSCs) of SGNs in 31 paired recordings. We stimulated the IHC with 10 ms depolarizations to different potentials ranging up to 57 mV in 5 mV steps (IV protocol; Fig. 3A). We deemed it incompatible with a reasonable productivity of the technically challenging, low-throughput paired recordings to combine them with imaging of Ca^2+^ at single AZs. Therefore, this study relies on analysis of the presynaptic Ca^2+^ influx at the level of the whole IHC (i.e. summing over all synapses and a low density of extrasynaptic Ca^2+^ channels, (Frank et al., 2009; Wong et al., 2014)). IHCs with synapses classified as high (*n* = 10) or low *SR* (*n* = 21) had similar Ca^2+^ current-voltage (IV) relationships: comparable maximal Ca^2+^ currents (Fig. 3Bi; *p =* 0.6939, Mann-Whitney U test) elicited at similar potentials (Fig. 3Bii; *p* = 0.1795, unpaired t-test) and comparable reversal potentials (Figure 3–figure supplement 1, panel A; *p* = 0.4034, unpaired t-test). The fractional activation of Ca^2+^ channels was determined from the normalized chord conductance of the IHC. Fitting a Boltzmann function to these activation curves (Fig. 3C, upper panel), we obtained the voltages of half-maximal activation (V_half Ca_) and the voltage-sensitivity of activation (slope) of the IHC Ca^2+^ channels.

“Release-stimulus intensity” curves, akin of an *ex vivo* representation of the SGN rate-level function, were constructed from the normalized Q_EPSC_ response obtained during the IV protocol (Figure 3–figure supplement 1, panel C). The voltage dependence of synaptic vesicle release per active zone was approximated by the fit of a sigmoidal function to the individual release-intensity curves (Fig. 3C, lower panel and Fig. 3Dii). From these sigmoidal fits, we obtained voltage of 10%- maximal release (Q_10, EPSC_), voltage of half-maximal release (Q_50, EPSC_), voltage of 90%-maximal release (Q_90, EPSC_) and the voltage sensitivity of release (slope). For two low *SR* paired recordings (Fig. 3Dii, grey dotted lines), a sigmoidal function did not properly fit the release-intensity curve (assessed by visual inspection) which led us to exclude them from the statistical analysis.

The voltage dependence of activation of whole-cell Ca^2+^ influx was similar between IHCs contacted by high and low *SR* boutons: threshold of Ca^2+^ influx (Fig. 3Ei; *p* = 0.2393, unpaired t-test), V_half Ca_ (Fig 3Fi; *p* = 0.3479, unpaired t-test) and voltage sensitivity of Ca^2+^ influx (Figure 3–figure supplement 1, panel B; *p* = 0.3470, unpaired t-test) did not differ significantly between IHCs contacted by low or high *SR* synapses. This seems to rule out a potential scenario in which the diverse SGN firing properties would be caused by varying average properties of Ca^2+^ channels among different presynaptic IHCs. Q_50, EPSC_ (Fig. 3Fii) of high *SR* synapses (−47.76 ± 1.4 mV; *n* = 10) was 6.76 ± 2.0 mV more negative compared to low *SR* synapses (−41.00 ± 1.2 mV; *n* = 19; *p* = 0.0021, unpaired t-test). Accordingly, high *SR* synapses had lower voltage thresholds of release than low *SR* synapses (Fig. 3Eii, Q_10, EPSC_ of −56.91 ± 1.5 mV [median −56.64 mV] vs. −47.39 ± 1.4 mV [median −48.89 mV]; *p* = 0.0001, Mann-Whitney U test). The hyperpolarized shift was not significant for Q_90 EPSC_ (Figure 3–figure supplement 1, panel D; *p* = 0.1706, unpaired t-test). The voltage sensitivity of release, determined by a slope factor, tended to be lower in high *SR* (4.16 ± 0.75 mV) than in low *SR* (2.9 ± 0.35 mV) synapses without reaching statistical significance (Figure 3–figure supplement 1, panel E; *p* = 0.0940, unpaired t-test;). The dynamic range, defined as the voltage range for which the exocytosis changes from 10-90% (Q_90 EPSC_ – Q_10 EPSC_), tended to be larger for high *SR* synapses without reaching significance (Fig. 3G; 18.30 ± 3.3 mV for high *SR* synapses vs 12.78 ± 1.5 mV for low *SR* synapses; *p* = 0.0940, unpaired t-test). The voltage dependence of activation of Ca^2+^ influx and of synaptic vesicle release did not differ significantly when the synapses where grouped based on their topographical location at the IHC (*n* = 12 modiolar synapses vs *n* = 17 pillar synapses; Figure 3–figure supplement 1, panels F-K). However, pillar *SR* synapses had a tendency to show lower voltage thresholds of release than modiolar synapses (Figure 3–figure supplement 1, panel Gii; Q_10, EPSC_ of −52.78 ± 1.8 mV [median −53.73 mV] vs. −47.68 ± 1.8 mV [median −49.30 mV]; *p* = 0.0725, Mann-Whitney U test). Altogether, these results demonstrate that high *SR* synapses release at more hyperpolarized voltages than low *SR* synapses.

Finally, we studied the apparent Ca^2+^ dependence of synaptic vesicle (SV) release during the aforementioned IV protocol, i.e., in the range of IHC receptor potentials. This protocol varies Ca^2+^ influx mainly via changing the channel open probability and to a lesser extent by changing the single channel current. We note that a supralinear intrinsic Ca^2+^ dependence of exocytosis in IHCs (i.e. Ca^2+^ cooperativity, *m ∼*3-4 when changing the single channel current) has been observed for IHCs of the cochlear apex in mice after hearing onset (Brandt, 2005; Jaime Tobón and Moser, 2023; Özçete and Moser, 2021; Wong et al., 2014). This is thought to reflect the cooperative binding of ∼4 Ca^2+^ ions required to trigger IHC exocytosis (Beutner et al., 2001). In contrast, a lower Ca^2+^ cooperativity was observed in these studies when primarily changing the number of open Ca^2+^ channels (*m < 2*). This difference in *m* observed for the apparent Ca^2+^ dependence of exocytosis has been taken to suggest a tight, Ca^2+^ nanodomain-like control of release sites by one or few Ca^2+^ channel(s) in line with classical studies of the Ca^2+^ dependence of transmitter release (Augustine et al., 1991). Here, we related changes of release at individual synapses (ΔQ_EPSC_) to the change of the integrated IHC Ca^2+^ influx (τιQ_Ca_). We fitted power functions (Q_EPSC_ = a(Q_Ca_)*^m^*) to the relationships for individual synapses (Figure 3–figure supplement 2) and found Ca^2+^ cooperativities of *m* < 2 for all but 2 synapses. This result suggests a tight, Ca^2+^ nanodomain-like control of release sites by one or few Ca^2+^ channel(s). Interestingly, however, high *SR* synapses, on average, had significantly lower Ca^2+^ cooperativities than low *SR* synapses (Fig. 3H; *m_highSR_* of 0.8 ± 0.1 [median 0.75; *n* = 10] vs *m_lowSR_* of 1.4 ± 0.1 [median 1.37; *n* = 21]; *p* = 0.0016, Mann-Whitney U test). The fit to pooled normalized data of high and low *SR* synapses yielded the same Ca^2+^ cooperativities of *m_highSR_* of 0.8 and *m_lowSR_* of 1.4 (Fig. 3I). When grouped based on their modiolar or pillar location, pillar synapses showed significantly lower Ca^2+^ cooperativities than modiolar synapses (Figure 3– figure supplement 1, panel L; *m_pillar_* of 1.0 ± 0.08 [median 0.88; *n* = 19] vs *m_modiolar_* of 1.6 ± 0.2 [median 1.3; *n* = 12]; *p* = 0.0202, Mann-Whitney U test). Our findings indicate that most afferent IHC synapses of hearing mice employ a tight, Ca^2+^ nanodomain-like control of release sites by one or few Ca^2+^ channel(s) for physiological sound encoding. Yet, quantitative differences in coupling seem to exist between high *SR*/pillar synapses and low *SR*/modiolar synapses, whereby a control of SV release by ∼1 Ca^2+^ channel prevails at high *SR*/pillar synapses.

### ii) Synaptic vesicle pool dynamics at individual IHC active zones

In 13 of the 31 aforementioned paired recordings (6 classified as low *SR* and 7 as high *SR*; 2 belonging to modiolar and 11 to pillar synapses), we employed a forward masking paradigm to study SV pool dynamics of single afferent synapses. The forward masking paradigm (Harris and Dallos, 1979) is commonly used for *in vivo* analysis of SGN spike rate adaptation and recovery from adaptation, which has been attributed to the depletion of readily releasable pool of SVs (RRP) and the recovery from depletion (Avissar et al., 2013; Frank et al., 2010; Furukawa and Matsuura, 1978; Goutman, 2017; Goutman and Glowatzki, 2007; Li et al., 2009; Moser and Beutner, 2000; Schroeder and Hall, 1974). Typically, the *in vivo* protocol is applied at saturating sound pressure levels, which we aimed to mimic using strong step IHC depolarizations (to −19 mV from −58 mV) separated by different interstimulus intervals (ISI: 4, 16, 64 and 256 ms) (Fig. 4A). In analogy to the *in vivo* forward masking paradigm, the first stimulus - called *masker*, as it depresses the response to a subsequent stimulus when applied in rapid succession- had a duration of 100 ms. The second stimulus -denominated *probe*- lasted for 15 ms. The recordings included a time frame of 400 ms preceding the “masker” and 400 ms following the “probe”, and the interval between masker and masker was 20 s. Applied to recordings of eEPSCs, the forward masking protocol provides experimental access to the initial RRP release rates, kinetics and extent of RRP depletion, sustained exocytosis, as well as recovery from RRP depletion. To accommodate the stochasticity of SV release from the RRP of IHC active zones, we run each protocol several times (≥3 to ≤ 20), which is routinely done for *in vivo* SGN physiology, but challenging *ex vivo* given the fragile and typically short-lived paired pre- and postsynaptic recordings (e.g. Goutman, 2017; Goutman and Glowatzki, 2007). Note that we did not employ cyclothiazide to inhibit AMPA receptor desensitization and reduce its contribution to postsynaptic eEPSC depression (Goutman, 2017), given the potential presynaptic effects of cyclothiazide in synaptic release (Diamond and Jahr, 1995; Dittman and Regehr, 1998).

**Figure 4.**
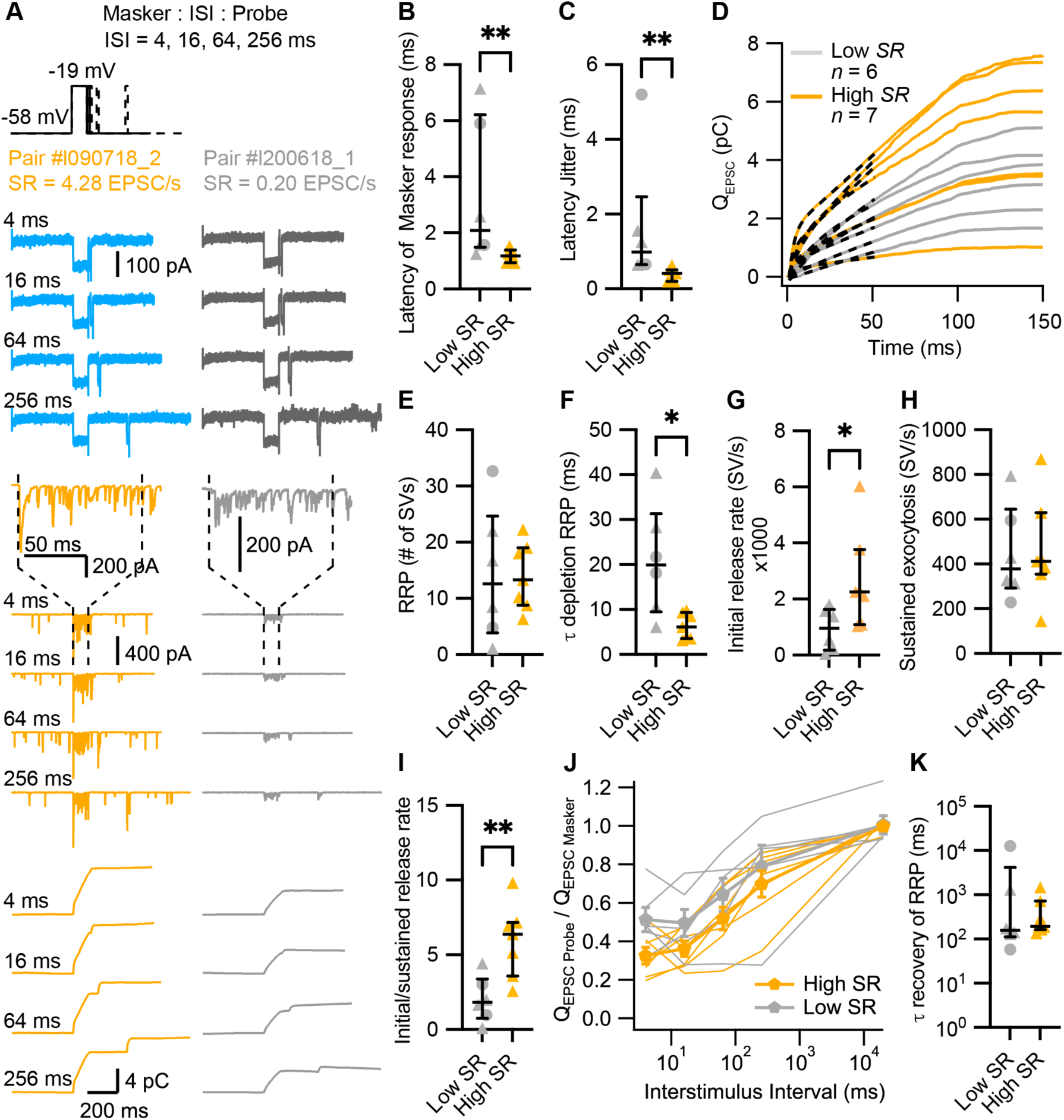
High *SR* synapses have shorter synaptic delay and higher initial release rates. **A.** “Forward Masking” voltage-protocol to study depletion and recovery of RRP and data of an exemplary high *SR* synapse (left panels, blue and orange) and low *SR* synapse (right panels, grey): Ca^2+^ currents (2^nd^ from top, I_Ca_), eEPSCs (2^nd^ from bottom) and Q_EPSC_ (bottom). The stimulus (top panel) consists of two sequential voltage steps (“masker” and “probe”) separated by different interstimulus intervals (ISI). Dashed vertical lines on top of eEPSC data indicate the onset and offset of the masker stimulus. **B.** Latencies of the evoked EPSCs (EPSC_onset_ - Masker_onset_) were significantly shorter in high *SR* than low *SR* synapses. **C.** High *SR* synapses also had less latency jitter. **D.** Pool depletion dynamics were studied by fitting the sum of a single exponential and a line function (black discontinuous line) to the first 50 ms of the average Q_EPSC_ trace in response to the masker stimulus. **E-I.** RRP, time constant (τ) of depletion, initial release rate and sustained release were calculated from the fits and the mean Q_sEPSC_ for each pair. High *SR* synapses depleted the RRP with faster time constants (F) and reached higher initial release rates (G) followed by a stronger adaptation (I). **J.** Recovery from RRP depletion shown as ratio of Q_EPSC probe_ and Q_EPSC masker_ (mean ± sem) during the first 10 ms of the stimulus. **K.** Time constant of recovery from RRP depletion obtained from single exponential fits to the traces shown in J (see Figure 4–figure supplement 1, panel L). Panels B, C, E-I and K show individual data points with the median and interquartile range overlaid (line). Synapses were classified as △ pillar or ❍ modiolar, and as Low *SR* < 1 sEPSC/s ≤ High *SR*.

For the analysis of evoked release dynamics, we focused on the response evoked by the masker. At the presynaptic level, there was no difference in the peak, initial and final IHC Ca^2+^ currents (I_Ca_) and Ca^2+^ current charge (Q_Ca_) between the recordings from high and low *SR* synapses (Figure 4–figure supplement 1, panel A-D). We calculated the synaptic delay from the onset of the masker to the onset of the eEPSC. High *SR* synapses had significantly shorter synaptic delays (Fig. 4B), with mean latencies of the first eEPSC after stimulus onset of 1.17 ± 0.09 ms (median 1.18) compared to 3.34 ± 1.04 ms (median 2.09) in low *SR* pairs (*p* = 0.0082, Mann-Whitney U test). This result was corroborated in a bigger sample size, when we compared the synaptic latencies of the 31 pairs to 10 ms pulses to −19 mV (Figure 4–figure supplement 1, panel E; latency of 1.19 ± 0.14 ms [median 1.14] in high *SR* compared to 2.57 ± 0.48 ms [median 1.79] in low *SR* synapses; *p* = 0.0101, Mann-Whitney U test). The latency jitter (measured as the standard deviation of the masker-evoked EPSC latencies; Fig. 4C) was also significantly smaller in high *SR* synapses compared to low *SR* synapses (0.38 ± 0.07 ms [median 0.41] vs 1.67 ± 0.72 ms [median 0.98], respectively; *p* = 0.0012, Mann-Whitney U test). Additionally, high *SR* synapses had significantly larger peak amplitudes of the masker-evoked EPSCs which reports the initial release from the RRP (Figure 4–figure supplement 1, panel F; −420.9 ± 54.22 pA for high *SR* synapses vs −240.1 ± 23.19 pA for low *SR* synapses; *p* = 0.0149, unpaired t-test). In contrast, the charge of masker-evoked EPSCs (Q_EPSC MASKER_) was more comparable (Figure 4–figure supplement 1, panel G; 4.512 ± 0.82 pC for high SR vs 3.020 ± 0.43 pC for low *SR* pairs; *p* = 0.1535, unpaired t-test). We fitted the first 50 ms of the Q_EPSC MASKER_ with the sum of a single exponential and a line function in order to analyze RRP depletion and sustained release (Figure 4–figure supplement 1, panel H; dashed lines in Fig. 4D). The amplitude of the exponential component *(A1)*, thought to reflect RRP exocytosis, was not different between high and low *SR* synapses (Figure 4–figure supplement 1, panel I; *p* = 0.4092, unpaired t-test). Likewise, the slope of the linear component, reflecting sustained exocytosis, did not differ significantly between the two groups (Figure 4–figure supplement 1, panel J; *p* = 0.1807, Mann-Whitney U test).

To quantify synaptic release in terms of SVs, we divided Q_EPSC MASKER_ by the mean Q_sEPSC_ recorded for each pair (see Fig. 2K). This builds on our assumption that each sEPSC corresponds to a unitary release event (“univesicular mode of release” (Chapochnikov et al., 2014; Grabner and Moser, 2018; Huang and Moser, 2018)). The quantal content (RRP size in #SV) was comparable between high and low *SR* synapses (Fig. 4E, 13.99 ± 2.239 SVs vs 14.28 ± 4.839 SVs, respectively; *p* = 0.9553, unpaired t-test). However, the RRP depleted significantly faster in high *SR* synapses (Fig. 4F): τ_RRP depletion_ was 6.347 ± 1.096 ms (median 6.13) for high *SR* vs. 20.88 ± 5.063 ms (median 19.93) for low *SR* synapses (*p* = 0.0140, Mann-Whitney U test). Accordingly, high *SR* synapses showed significantly higher initial release rates compared to low *SR* synapses (Fig. 4G; 2651 ± 660.0 SV/s vs. 927.4 ± 307.4 SV/s; *p* = 0.0472, unpaired t-test), which given the comparable RRP size, indicates a higher release probability of the high *SR* synapses. Moreover, high *SR* synapses showed stronger depression of the release rate (Fig. 4I; the ratio of initial/sustained release rate was 5.941 ± 0.916 for high *SR* pairs and 2.023 ± 0.6251 for low *SR* pairs; *p* = 0.006, unpaired t-test), despite comparable sustained release rates (Fig. 4H; *p* = 0.9258, unpaired t-test).

Finally, we determined RRP recovery from depletion using the ratio Q_EPSC Probe_/Q_EPSC Masker_, whereby Q_EPSC_ for masker and probe was estimated for the first 10 ms of stimulation. Figure 4J plots the ratio for each interstimulus interval (ISI), including the masker-to-masker interval. All the synapses exhibited a smaller response (Fig. 4J) and a longer response latency (Figure 4–figure supplement 1, panel K) to the probe compared to the response to the masker stimulus. This short-term synaptic depression probably reflects RRP depletion by the masker stimulus (Avissar et al., 2013; Cho et al., 2011; Frank et al., 2010; Moser and Beutner, 2000), in contrast to previous reports of synaptic facilitation when employing short paired stimuli from more hyperpolarized resting potential that do not trigger the full RRP release (Chen and von Gersdorff, 2019; Goutman and Glowatzki, 2011). Surprisingly, and contrary to a previous report in auditory bullfrog synapses (Cho et al., 2011), approximately half of the synapses showed a lower ratio at 16 ms than at 4 ms regardless of their *SR*. Therefore, to determine the kinetics of recovery from RRP depletion, we fitted a single exponential to the recovery data over the ISI range from 16 ms to 20 s (Figure 4-figure supplement 1, panel L). We did not find a significant difference between high *SR* and low *SR* synapses (Fig. 4K; τ_recovery RRP_: 451.5 ± 187.0 ms [median 192.7] vs 2416 ± 2064 ms [median 157.9], respectively; *p* = 0.5338, Mann-Whitney U test). In 4 high *SR* and 4 low *SR* synapses, spontaneous activity was resumed shortly after the offset of the probe. The time to this first sEPSC took longer in high *SR* synapses compared to low ones (Figure 4–figure supplement 1, panel M; 185.2 ± 25.25 ms [median 161.6] vs 104.5 ± 24.10 ms [median 108.4], respectively; *p* = 0.0286, Mann-Whitney U test), again indicating stronger synaptic depression on high *SR* synapses.

## Discussion

Much of the information on synaptic sound encoding at afferent IHC-SGN synapses has been obtained from either juxtacellular recordings of SGN firing *in vivo* or from *ex vivo* patch-clamp recordings. Yet, it has remained difficult to reconcile those *in vivo* and *ex vivo* results and to establish a unified account of sound intensity coding in the auditory nerve given differences in experimental conditions, animal models and protocols employed. Here, we biophysically characterized the heterogeneous function of afferent SGN synapses in hearing mice with reference to their rate of spontaneous transmission (*SR*) as a surrogate of SGN SR that informs their functional properties. We performed paired pre- and post-synaptic patch-clamp recordings from single IHC synapses of hearing mice under near-physiological conditions using protocols adapted from *in vivo* characterization of SGN’s response properties. Using this approach, we were able to distinguish synapses with low and high *SR*, which we propose to provide the input into low and high SR SGNs. We found that about 90% of high *SR* synapses were located at the pillar side of the IHC. High *SR* synapses had larger sEPSCs with a monophasic (or more compact) waveform, lower voltage-thresholds of release, shorter synaptic delays, tighter coupling of release sites to Ca^2+^ channels as well as higher initial release rates and shorter RRP depletion time constants. RRP size, rate of sustained exocytosis and kinetics of RRP recovery from depletion were comparable between high and low *SR* synapses. We conclude that high *SR* synapses exhibit higher release probability which likely reflects the tighter coupling of Ca^2+^ channels and release sites. This diversity in the response properties of individual synapses most likely expands the capacity of a single IHC to encode sound intensity over the wide range of audible sound pressures.

### Diversity of spontaneous release and their topographical segregation

The *SR* range observed in our paired recordings from mouse afferent synapses (0 to 18 EPSC/s) agrees with results obtained without patch-clamping the IHC (0 to 16 sEPSC/s) and with previous *ex vivo* reports using loose patch recordings from SGNs of p15-p17 rats (0.1 to 16.42 spikes/s; (Wu et al., 2016). However, the maximum rate is considerably smaller than those recorded *in vivo* from single SGNs of p14-p15 mice (up to 60 spikes/s; Wong et al., 2013). This 3-fold difference between *ex vivo* and *in vivo* recordings of the same age group could indicate that *in vivo*, the IHC resting potential might be more depolarized and/or subject to spontaneous fluctuations that can trigger Ca^2+^ channel openings and release. Additionally, the presence of K^+^ channel blockers (TEA and Cs^+^) and differences in pH can all have an impact on the excitability of the cell and the kinetics of the cellular processes.

Other important factors might explain the higher SR reported *in vivo*: i) The intrinsic biophysical properties of SGNs which could further expand the firing rate distribution (Markowitz and Kalluri, 2020), for instance, due to spikes initiated intrinsically and not activated by an EPSP (Wu et al., 2016); ii) A sampling bias of synapses with *SR* above 20 sEPSC/s given that they constitute less than 35% of SGNs from 8-17 weeks old mice (Taberner and Liberman, 2005); and, iii) the developmental recruitment of high SR SGNs with age (Niwa et al., 2021; Romand, 1984; Walsh and McGee, 1987; Wong et al., 2013; Wu et al., 2016). 90% of the paired recordings (and 60% for the bouton recordings) of our dataset were obtained from mice between p14-p17, in which spontaneous activity is still low compared to older age groups (p19-p21: 0 – 44.22 spikes/s; p29- p32: 0.11 – 54.9 spikes/s (Wu et al., 2016); p28: 0 – 47.94 spikes/s, (Siebald et al., 2023)).

In analogy to the pioneering finding of a synaptic segregation of cat SGNs according to SR along the pillar–modiolar axis of IHCs (Liberman, 1982; Merchan-Perez and Liberman, 1996), we found that about 90% of high *SR* were located on the pillar IHC side. Yet, not all the synapses of the pillar IHC side had high *SR*, which agrees with a recent study of molecularly tagged SGNs (Siebald et al., 2023). These findings suggest that high frequency and large amplitude sEPSCs occur predominantly in synapses with smaller ribbons and AZs, opposing results from retinal cells in which smaller ribbons resulted in reduced frequency and amplitude of EPSCs (Mehta et al., 2013). It is important to point out that our modiolar/pillar classification is less precise than that of other studies in which the synapse position was quantitatively assigned (Frank et al., 2009; Liberman et al., 2011; Ohn et al., 2016; Özçete and Moser, 2021). Moreover, other studies also support an overall pillar-modiolar gradient with “salt and pepper” intermingling of synaptic properties rather than their strict segregation (Ohn et al., 2016; Özçete and Moser, 2021).

Similar to previous reports (e.g. Chapochnikov et al., 2014; e.g. Glowatzki and Fuchs, 2002; Grant et al., 2010; Huang and Moser, 2018; Rutherford et al., 2012), we found a high variability in the waveform and amplitude of sEPSCs between synapses, which apparently do not strictly depend on the topographical location of the synapse (the present work and (Niwa et al., 2021)). Yet, high *SR* synapses had larger and monophasic (or temporally more compact) sEPSCs. This likely also explains the shorter 10-90% rise time of the sEPSC in high *SR* synapses, as monophasic sEPSC have shorter rise times than the multiphasic ones (Chapochnikov et al., 2014; Glowatzki and Fuchs, 2002; Huang and Moser, 2018). The difference in the percentage of monophasic sEPSCs and rise times in low and high *SR* synapses could arise from variability in the fusion pore dynamics on the way to SV fusion and/or on the number of SVs released in timely manner. In the view of the multivesicular hypothesis of spontaneous release (Glowatzki and Fuchs, 2002; Li et al., 2009; Niwa et al., 2021; Schnee et al., 2013), SVs of a modiolar AZ might fuse in an uncoordinated manner, creating EPSCs with a less compact waveform and slower rise times.

Alternatively and our favorite hypothesis, each sEPSC corresponds to a unitary release event (“univesicular mode of release” (Chapochnikov et al., 2014; Grabner and Moser, 2018; Huang and Moser, 2018; Young et al., 2021)) that has the capacity to drive action potential firing (Rutherford et al., 2012). In the framework of the univesicular hypothesis of spontaneous release, the flickering of the fusion pore, prior to or instead of full collapse fusion of the SV, might be favored in low *SR* synapses, leading to slower sEPSC rise times and a lower percentage of monophasic sEPSCs. Such heterogeneity of fusion pore dynamics has been reported in chromaffin cells, calyx of Held and hippocampal neurons (Chang et al., 2021; Henkel et al., 2019; Shin et al., 2020, 2018).

Finally, we note that sEPSC amplitudes of IHC synapses in hearing mice (present study and (Niwa et al., 2021)) seem lower than in previous *ex vivo* studies on IHC synapses of hearing rats (Chapochnikov et al., 2014; Grant et al., 2010; Huang and Moser, 2018; Young et al., 2021). In rats, the EPSC amplitude distribution changes with maturation, from highly skewed to the left with a peak around −30 pA to a Gaussian-like distribution with a peak at −375 pA (Grant et al., 2010). This does not seem to be the case in mouse IHC synapses. Average EPSC amplitudes in pre-hearing mice are around −100 to −150 pA (Chapochnikov et al., 2014), even with 40 mM K^+^ stimulation (Jing et al., 2013; Takago et al., 2019). On the contrary, mean EPSC amplitudes in hearing mice remained small (around −100 pA) in resting conditions ((Niwa et al., 2021) and the present study), but became significantly larger upon stimulation with 40 mM K^+^ (Niwa et al., 2021) or voltage depolarizations (the present study, Figure 1-figure supplement 1, panel G).

### Candidate mechanisms distinguishing evoked release at low and high SR synapses

The temporal and quantal resolution offered by paired recordings allowed us to analyze the biophysical properties of evoked synaptic transmission in relation to the *SR* of the given synapse. In an intriguing resemblance with *in vivo* evoked firing properties of high SR SGNs (Bourien et al., 2014; Buran et al., 2010; Relkin and Doucet, 1991; Rhode and Smith, 1985; Taberner and Liberman, 2005), high *SR* synapses showed lower voltage (∼sound pressure *in vivo*) thresholds of synaptic transmission (∼firing *in vivo*), shorter and less variable synaptic latencies (∼first spike latencies *in vivo*), and higher initial release rates (∼onset firing rate *in vivo*). In addition, we found stronger synaptic depression at high *SR* synapses, which agrees well with the finding of a greater ratio of peak to adapted firing rate in high SR SGNs recorded *in vivo* (Taberner and Liberman, 2005). These results support the hypothesis that IHC synaptic heterogeneity (Frank et al., 2009; Hua et al., 2021; Ohn et al., 2016; Özçete and Moser, 2021; Reijntjes et al., 2020) contributes to the diversity of spontaneous and sound-evoked SGN firing.

How do high *SR* synapses with likely smaller ribbons and lower maximal Ca^2+^ influx achieve a shorter latency and higher initial release rate? Our hypothesis is that, in high *SR* synapses, a more hyperpolarized Ca^2+^ channel activation (Ohn et al., 2016) in combination with tighter coupling between the Ca^2+^ channels and the Ca^2+^ sensor of fusion (this work and (Özçete and Moser, 2021)) would enable a faster response with a greater initial SV release probability for a given stimulus. The spatial coupling of the Ca^2+^ channel to the SV release site has also been shown to greatly affect release probability in other synapses (Eggermann et al., 2012; Fekete et al., 2019; Moser et al., 2019; Rebola et al., 2019). Thus, a heterogenous Ca^2+^ coupling would diversify the response properties (i.e., synaptic vesicle release probability) of individual synapses to the same stimulus. In IHCs, this is particularly important for sound intensity and temporal coding (reviewed in (Moser et al., 2023)). Interestingly, genetic disruptions that shifted the voltage-dependence had a greater impact on the *in vivo* distribution of SR and onset firing rate of SGNs than mutations that changed the maximal synaptic Ca^2+^ influx (Jean et al., 2018; Ohn et al., 2016).

Paired pre- and postsynaptic patch-clamp recordings (this work) and single synapse imaging of presynaptic Ca^2+^ signals and glutamate release (Özçete and Moser, 2021) jointly found a lower apparent Ca^2+^ cooperativity in pillar synapses during depolarizations within the range of receptor potentials. The sensitivity and temporal resolution of paired recordings further allowed us to classify the synapses based on their spontaneous rate and support the hypothesis that high SR SGNs receive input from AZ with tighter coupling than low SR SGNs. However, single synapse imaging (Özçete and Moser, 2021) found a wider range of apparent Ca^2+^ cooperativities than our two non-overlapping data sets for paired patch-clamp recordings (this work and (Jaime Tobón and Moser, 2023)). This might reflect two important technical differences: i) single synapse imaging assessed the presynaptic Ca^2+^ influx of the specific synapse, while in paired recordings we related release to the whole cell Ca^2+^ influx, and ii) the temporal resolution of paired recordings allowed to study the initial release rate using shorter stimuli than in imaging, which avoids an impact of RRP depletion and ongoing SV replenishment. Future studies, potentially combining paired patch-clamp recordings with imaging of presynaptic Ca^2+^ signals, will be needed to further scrutinize the heterogeneity of Ca^2+^ dependence of release in IHCs and its impact on release probability.

Other factors that affect release probability include variations in the number of open Ca^2+^ channels at the AZ (Gratz et al., 2019; Holderith et al., 2012; Scimemi and Diamond, 2012; Sheng et al., 2012; Wong et al., 2013) and the fusion competence of the SV (Klenchin and Martin, 2000), including the priming and docking state (Lin et al., 2022; Neher and Brose, 2018). The ensuing Ca^2+^ influx in pillar synapses might facilitate Ca^2+^ channels and priming of SVs (Cho and von Gersdorff, 2012; Goutman and Glowatzki, 2011, 2007; Michalski et al., 2017; Moser and Beutner, 2000; Pangrsic et al., 2015; Schnee et al., 2011; Spassova et al., 2004) and contribute to the observed results in high *SR* synapses. Regarding the fusion competence of SVs, it is unknown whether modiolar and pillar synapses exhibit different numbers of docked and primed SVs. To date, ultrastructural studies that resolve docked and tethered SVs have not addressed the topographical location of the AZ in the murine IHC (Chakrabarti et al., 2022, 2018).

Besides SV release probability, RRP size co-determines neurotransmitter release. Our estimated RRP of about 14 SVs in both high and low *SR* synapses compares well to prior estimates obtained using *ex vivo* electrophysiology (10-40 SVs: Refs. (Goutman and Glowatzki, 2007; Jean et al., 2018; Johnson et al., 2005; Khimich et al., 2005; Moser and Beutner, 2000; Pangrsic et al., 2010; Schnee et al., 2005), model-based analysis of SGN firing (4-40 SVs (Frank et al., 2010; Jean et al., 2018; Peterson et al., 2014)) and electron microscopy (10-16 SVs within 50 nm of the presynaptic membrane (Chakrabarti et al., 2018; Frank et al., 2010; Graydon et al., 2011; Kantardzhieva et al., 2013; Khimich et al., 2005; Pangrsic et al., 2010)). However, previous reports based on electron microscopy (Kantardzhieva et al., 2013; Merchan-Perez and Liberman, 1996; Michanski et al., 2019) suggested larger pools of SVs at modiolar synapses, while our electrophysiological estimate of RRP size was comparable between low and high *SR* synapses. This finding argues against a strong contribution of RRP size to the observed differences in neurotransmitter release. However, higher release probability with comparable RRP size explains higher initial release rates, which likely explain the faster and temporally more precise postsynaptic depolarization that is likely to turn into shorter first spike latencies and lower first spike latency jitter (this study and (Buran 2010).

Finally, the heterogeneity in the functional properties of IHC synapses could arise from molecular heterogeneity of the AZ. In central glutamatergic synapses, molecular heterogeneity of synaptic proteins plays a critical role in the modulation of SV release probability and priming state (Neher and Brose, 2018; Wichmann and Kuner, 2022). For instance, differential isoforms of priming factors and scaffold proteins have been suggested to tune the functional synaptic diversity of central synapses (Fulterer et al., 2018; Rebola et al., 2019; Rosenmund et al., 2002). Cochlear IHCs have an unconventional fusion machinery that appears to work without neuronal SNARES (Nouvian et al., 2011) (but see Calvet et al., 2022) and priming factors such as Munc13 and CAPS (Vogl et al., 2015). Therefore, future studies will need to determine the molecular nanoanatomy underlying the specific AZ nanophysiology and functional synaptic heterogeneity at IHCs. Promising candidates include RBPs (Butola et al., 2021; Grauel et al., 2016; Krinner et al., 2021, 2017; Petzoldt et al., 2020), RIMs (Jung et al., 2015; Maria M. Picher et al., 2017), and Septin (Fekete et al., 2019; Yang et al., 2010).

### Challenges for relating synaptic and neural response properties

Next to providing support for the presynaptic hypothesis of functional SGN diversity, the present study also highlights some of the challenges met when aiming to bridge the gap between presynaptic hair cell function and neural sound encoding. Despite major efforts undertaken to match experimental conditions and protocols, it remains difficult to reconcile some findings of *ex vivo* and *in vivo* physiology. Parameters such as RRP size (∼# spikes of the rapidly adapting component of firing), sustained exocytosis (∼adapted firing rate *in vivo*), recovery of spontaneous and evoked release (∼recovery from forward masking *in vivo*) did not differ among our high and low *SR* synapses, and contrasts with *in vivo* data (e.g. Refs. (Bourien et al., 2014; Buran et al., 2010; Relkin and Doucet, 1991; Rhode and Smith, 1985; Taberner and Liberman, 2005)).

Of particular interest is that the dynamic ranges and slope of release-intensity relationship of high and low *SR* synapses diverge from the expectations if assuming that high SR SGNs are driven by high *SR* synapses. High *SR* synapses tended to show broader dynamic ranges with shallower slopes, while, *in vivo,* high SR SGNs show smaller dynamic ranges and steeper slopes than the low SR ones (Ohlemiller et al., 1991; Winter et al., 1990). Could this reflect the non-linear saturating properties of the basilar membrane (Sachs et al., 1989; Sachs and Abbas, 1974; Yates et al., 1990) (discussed in Ref. (Ohlemiller et al., 1991)) which might widen the rate level function of low SR SGNs? Or is it due to a partial depletion of the “standing” RRP (i.e. the occupancy of the RRP release sites with a fusion-competent SV (Moser, 2020; Pangrsic et al., 2010)) at high *SR* synapses *in vivo*? It remains to be determined whether this and the other aforementioned differences between our data and *in vivo* reports could be attributed to mechanisms downstream of glutamate release and AMPA receptor activation. The possible mechanisms include but are not limited to: i) different spike rates due to diverse EPSC waveforms (Rutherford et al., 2012); ii) differences in SGN excitability (Crozier and Davis, 2014; Markowitz and Kalluri, 2020; Smith et al., 2015) due to heterogenous molecular (Petitpré et al., 2020, 2018; Shrestha et al., 2018; Sun et al., 2018) and morphological profiles (Liberman, 1980; Merchan-Perez and Liberman, 1996; Tsuji and Liberman, 1997); and iii) differences in efferent innervation of SGNs (Hua et al., 2021; Liberman, 1990; Ruel et al., 2001; Wu et al., 2020; Yin et al., 2014). Certainly, caution is to be applied for the comparison of *ex vivo* and *in vivo* data due to the partial disruption of the physiological milieu despite our efforts to maintain near-physiological conditions, and the incomplete synaptic maturation when focusing *ex vivo* experiments on the 3^rd^ postnatal week soon after hearing onset.

Clearly more work is needed to elucidate the mechanisms of SGN firing diversity in the cochlea. Ideally, future studies will combine *in vivo* and *ex vivo* experiments, such as combining physiological SGN characterization with neural backtracing and synaptic morphology of labelled SGNs using volume imaging of afferent and efferent connectivity (Hua et al., 2021). Moreover, combining optogenetic IHC stimulation with imaging of SGN activity could provide higher throughput and serve posthoc morphology. Finally, paired patch-clamp recordings, as done in the present study, could be combined with SGN subtype-specific molecular labeling, fiber tracing and immunolabeling to further relate synaptic transmission and SGN neurophysiology.

## Materials and Methods

### Animals and tissue preparation

c57BL/6N mice of either sex between postnatal day 14-27 were used. For paired recordings, the number of animals per age was: p14 (9), p15 (7), p16 (9), p17 (5), p18 (2), p20 (1). For the bouton recordings of Figure 2–figure supplement 1, panel A, the number of animals per age was: p14 (2), p15 (3), p16 (3), p21 (1), p24 (1), p25 (1), p27 (1). The animal handling and experiments complied with national animal care guidelines and were approved by the University of Göttingen Board for animal welfare and the Animal Welfare Office of the State of Lower Saxony. Animals were sacrificed by decapitation and the cochleae were extracted in modified Hepes Hank’s solution containing: 5.36 mM KCl, 141.7 mM NaCl, 1 mM MgCl_2_-6H_2_O, 0.5 mM MgSO_4_-7H_2_O, 10 mM HEPES, 0.5 mg/ml L-glutamine, and 1 mg/ml D-glucose (pH 7.2, osmolarity of ∼300 mOsm). The apical coil of the organ of Corti was dissected and placed under a grid in the recording chamber. Pillar or modiolar supporting cells were removed using soda glass pipettes in order to gain access to the basolateral face of the IHCs and to the postsynaptic boutons of type I SGNs. Dissection of the organ of Corti and cleaning of the supporting cells were performed at room temperature (20-25°C).

### Electrophysiological recordings

Pre- and postsynaptic paired patch clamp recordings were performed at near physiological temperature (32-37°C) using an EPC-9 amplifier (HEKA electronics) (Fig. 1). Patch electrodes were positioned using a PatchStar micromanipulator (Scientifica, UK). Whole-cell recordings from IHCs were achieved using the perforated-patch clamp technique (Moser and Beutner, 2000) using Sylgard™–coated 1.5 mm borosilicate pipettes with typical resistances between 3.5 and 6 MΩ. The IHC pipette solution contained: 129 mM Cs-gluconate, 10 mM tetraethylammonium (TEA)-Cl, 10 mM 4-AP, 10 mM HEPES, 1 mM MgCl_2_ (pH 7.2, osmolarity of ∼290 mOsm), as well as 300 μg/ml amphotericin B added prior to the experiment. Once the series resistance of the IHC reached below 30 MΩ, whole-cell voltage-clamp recordings from a contacting bouton was established as described in previous studies (Glowatzki and Fuchs, 2002; Grant et al., 2011; Huang and Moser, 2018). For 2 pairs, the bouton recording was established first and then the IHC. Sylgard™-coated 1.0 mm borosilicate pipettes with typical resistances between 7 and 12 MΩ were used for the postsynaptic recordings. The bouton pipette solution contained: 137 mM KCl, 5 mM EGTA, 5 mM HEPES, 1 Mm Na_2_-GTP, 2.5 mM Na_2_-ATP, 3.5 mM MgCl_2_·6H_2_O and 0.1 mM CaCl_2_ (pH 7.2 and osmolarity of ∼290 mOsm). The organ of Corti was continuously perfused with an extracellular solution containing 4.2 mM KCl, 95-100 mM NaCl, 25 mM NaHCO_3_, 30 mM TEA-Cl, 1mM Na- Pyruvate, 0.7 mM NH_2_PO_4_·H_2_O, 1mM CsCl, 1 mM MgCl_2_·H_2_O, 1.3 mM CaCl_2_, and 11.1 mM D-glucose (pH 7.3, osmolarity of ∼310 mOsm). 2.5 µM tetrodotoxin (Tocris or Santa Cruz) was added to block voltage-gated Na^+^ channels in the postsynaptic bouton.

Data were acquired using the Patchmaster software (HEKA electronics). The current signal was filtered at 5-10 kHz and sampled at 20-50 kHz. IHC were voltage-clamped at a holding potential of −58 mV, around the presumed *in vivo* resting potential (Johnson, 2015). The bouton was held at a potential of −94 mV. All reported potentials were corrected for the liquid junction potential (19 mV for the IHC and 4 mV for the bouton), measured experimentally. Ca^2+^ current recordings were corrected for the linear leak current using a *P/n* protocol. We excluded IHCs and boutons with leak currents exceeding −60 pA and −100 pA at holding potential, respectively. Average IHC series resistance (R_s_) was 14.7 ± 0.8 MΩ (14.63 ± 1.04 MΩ for high *SR* synapses vs 14.83 ± 1.07 MΩ for low *SR*; *p* = 0.7433, Mann-Whitney U test). Average IHC membrane capacitance (C_m_) was 8.8 ± 0.19 pF (8.8 ± 0.25 pF for recordings of high *SR* synapses vs. 8.8 ± 0.25 pF for low *SR*; *p* = 0.6237, Mann-Whitney U test). The apparent series resistance of the bouton was calculated from the capacitive transient in response to a 10-mV test pulse. The actual R_s_ was offline calculated as reported in (Huang and Moser, 2018). Briefly, we fitted the decay phase of the capacitive transient with a double exponential. Average bouton R_s_ from paired recordings was 64.6 ± 3.3 MΩ (60.2 ± 5.3 MΩ for high *SR* synapses vs 57.8 ± 3.6 MΩ for low *SR*; *p* = 0.7115, unpaired t-test; and 57.01 ± 4.7 MΩ for bouton recordings only). [Average R_s_ from the bouton recordings from Figure 2–figure supplement 1 was 57 ± 4.7 MΩ]. Bouton capacitance was estimated from the area under the fast component of the double exponential fit. Average bouton C_m_ was 1.7 ± 0.09 pF (1.8 ± 0.19 pF for high *SR* synapses vs 1.7 ± 0.11 pF for low *SR*; *p* = 0.6575, unpaired t-test). Average bouton membrane resistance (R_m_) was 1491 ± 133.2 MΩ (1499 ± 193.3 MΩ for high *SR* synapses vs 1487 ± 174.3 MΩ for low *SR*; *p* = 0.7143, Mann-Whitney U test). The properties of the recordings (i.e., amplitude of EPSC) were not correlated with the bouton R_s_ (Figure 1–figure supplement 1).

The threshold for sEPSC detection was 4 times SD of the baseline. Spontaneous activity was calculated from time windows without stimulation with the IHC held at −58 mV; either from a 5 – 10 s recording or by averaging the number of events from the segments before and after a depolarizing pulse (Fig. 1B, Supplementary Table 1). To study the depletion and recovery of the pool of vesicles, we used a protocol adapted from the forward masking protocol performed during *in vivo* extracellular recordings of SGNs (Harris and Dallos, 1979; Jean et al., 2018). It consisted of two consecutive depolarizing pulses to the voltage that elicited the highest peak of Ca^2+^ current (−19 mV; Fig. 1C). The first pulse, called masker, lasted 100 ms and it was followed by a second pulse, called probe, which lasted 15 ms. The two pulses were separated by intervals without depolarization (interstimulus intervals, ISI) that lasted 4, 16, 64 and 256 ms. The waiting time between masker and masker was 20 s and each protocol was repeated between 3 – 20 times. To study the dynamic voltage range of synaptic transmission, we used a current-voltage (IV) protocol with 10 ms pulses of increasing voltage (from −70 mV/-60 mV to 70 mV in 5 mV steps). The interval between two stimuli was 1.5 s.

### Data Analysis

Electrophysiological data was analyzed using the IgorPro 6 Software Package (Wavemetrics), GraphPad Prism 9 and Excel. Ca^2+^ charge (Q_Ca_) and EPSC charge (Q_EPSC_) were estimated by taking the integral of the current. Kinetics of sEPSCs, such as amplitude, 10-90% rise time, time constant of decay (τ_decay_) and full-width half-maximum (FWHM), were calculated with Neuromatic (Rothman and Silver, 2018).

To obtain IV curves, we averaged the evoked Ca^2+^-currents (I_Ca_) during 5 to 10 ms after the start of each depolarization. Fractional activation of the Ca^2+^ channels was obtained from the normalized chord conductance, *g*,

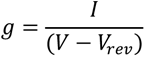

where V is the membrane potential and V_rev_ is the reversal potential determined by fitting a line function between the voltage of I_Ca peak_ + 10 mV and the maximal depolarization. The activation curve was approximated by a first-order Boltzmann equation:

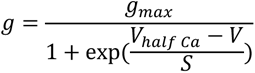

where *g_max_* is the maximum chord conductance, V_half Ca_ is the membrane potential at which the conductance is half activated, and *S* is the slope factor describing the voltage sensitivity of activation.

Release intensity curves were obtained by calculating Q_EPSC_ by the end of each depolarization step and fitted using a sigmoidal function:

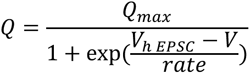

where Q_max_ is the maximal Q_EPSC_ (normalized to 1), V_h EPSC_ corresponds to the voltage of half-maximal release (or Q_50, EPSC_) and Q is the EPSC charge. The dynamic range was determined as the voltage range between 10% and 90% of the maximal vesicle release. For statistical analysis of dynamic range, we included only pairs for which both the Ca^2+^ fractional activation and the rate level curves were possible to fit.

The apparent Ca^2+^ dependence of neurotransmitter release was studied from the 10 ms step- depolarizations of the IV curves. The resulting Q_EPSC_ vs IHC Q_Ca_ plots from each individual pair were fitted with a power function:

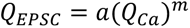

where *m* corresponds to the Ca^2+^ cooperativity. Some pairs showed a clear saturation of release at high IHC Q_Ca_. In these cases, the fit was restricted to the datapoints before the plateau, which was determined by visual inspection. For the pooled data, the power function was fitted to the normalized Q_EPSC_ vs normalized Q_Ca_. For the pairs with saturation of release, Q_Ca_ was normalized to a point before the plateau.

For forward masking experiments, the postsynaptic response was averaged for all the repetitions for each paired recording (between 3 and 20, depending on the stability of the pair). Single active zone pool dynamics were determined by fitting an exponential plus line function to the to the first 50 ms of the average Q_EPSC_ trace in response to the masker stimulus for each ISI,

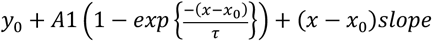

where *A1* is the amplitude of the exponential component, τ is the time constant of the exponential component. RRP size (in synaptic vesicles) was estimated from dividing *A1* by the charge of the average sEPSC for each pair. Sustained exocytosis rate (in SV per s) was calculated from the slope of the linear function divided the charge of the average sEPSC. Individual recovery kinetics were determined from the ratio of probe and masker responses at 10 ms of the depolarization, with the ratio between masker and masker being 1. The recovery traces were fitted with a single exponential function from 16 ms to 20 000 ms to determine the time constant of RRP recovery.

Data was prepared for presentation using Adobe Illustrator. Skewness analysis, PCA and other statistical analysis were performed using GraphPad Prism 9. Statistical significance was assessed with unpaired t-test or non-parametric Mann-Whitney U test depending on the normal distribution and equality of variances of the data (Saphiro-Wilk test and F test). Data is expressed as mean ± sem.

## Author Contributions

TM and LMJT designed the study. LMJT performed the experiments and the analysis. TM and LMJT prepared the manuscript.

## Acknowledgments

We would like to thank Dr. Chao-Hua Huang and Dr. Jakob Neef for their experimental and analytical input. Dr. Antoine Huet for the discussion regarding *in vivo* response properties of spiral ganglion neurons. Dres. Erwin Neher, Manfred Lindau and Jakob Neef for critical input into this project. LMJT was a recipient of the Erwin Neher Fellowship and T.M. is a Max-Planck Fellow at the Max Planck Institute for Multidisciplinary Sciences. This work was further supported by the Deutsche Forschungsgemeinschaft (DFG, German Research Foundation) via the Collaborative Research Center 889 (project A02) and the Leibniz Program (MO896/5 to T.M), the European Research Council through the Advanced Grant ‘DynaHear” to T.M. under the European Union’s Horizon 2020 Research and Innovation program (grant agreement No. 101054467), and by Fondation Pour l’Audition (FPA RD-2020-10). LMJT is a member of the Hertha Sponer College from the Cluster of Excellence Multiscale Bioimaging (MBExC). Open access funding provided by Max Planck Society.

---

**Figure 1–figure supplement 1.**
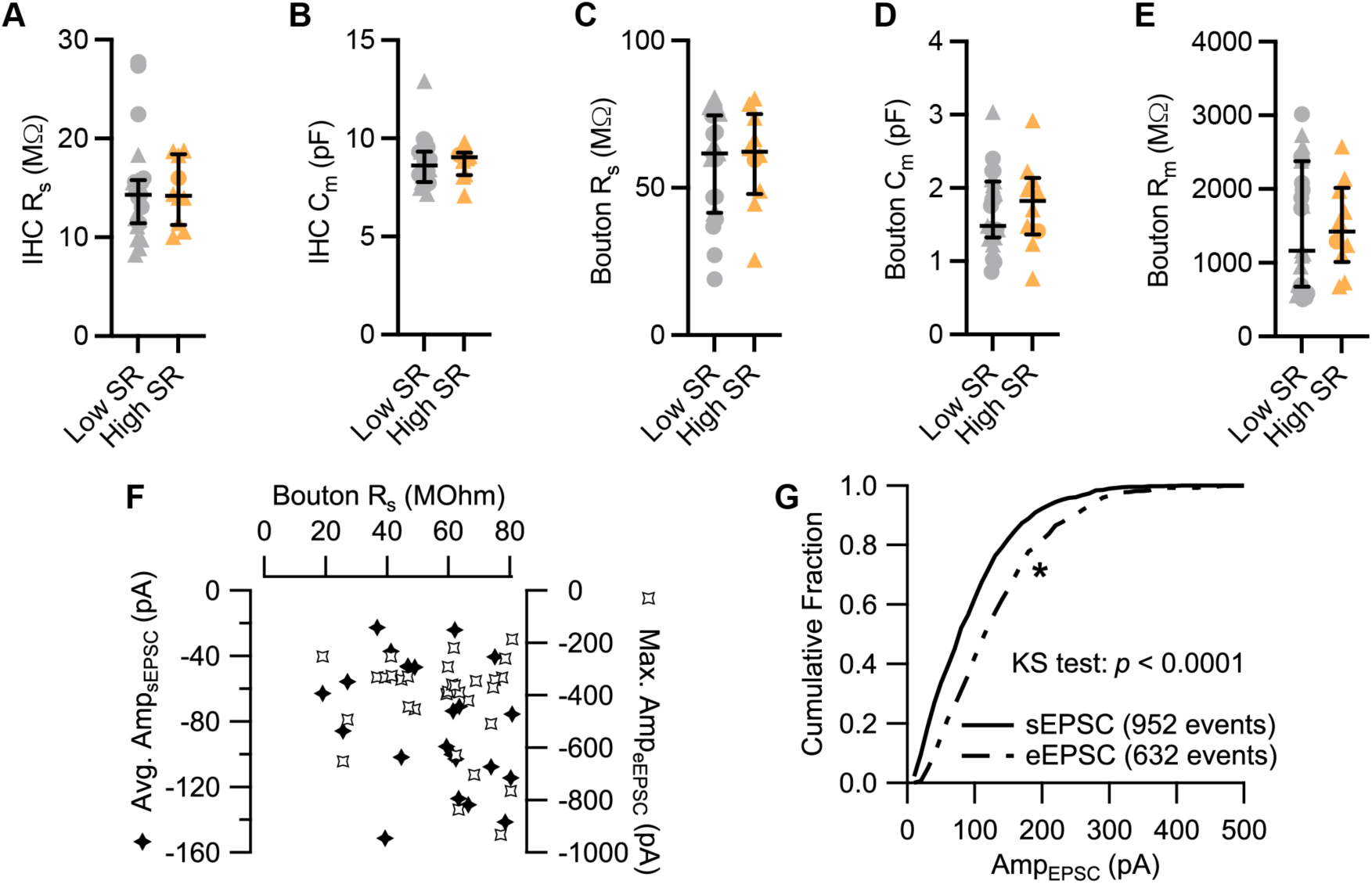
Passive electrical properties of IHCs and SGN boutons. **A-E.** Values for IHC’s series resistance (R_s_, A) and membrane capacitance (C_m_, B), and bouton’s R_s_ (C), C_m_ (D) and membrane resistance (R_m_, E) were comparable between paired recordings with low and high *SR*. **F.** Average sEPSC amplitudes and maximal eEPSC amplitudes plotted against bouton R_s_. **G.** Cumulative fraction of amplitude of sEPSCs and of eEPSCs recorded from the 13 synapses included in Figure 4. The eEPSCs correspond to voltage-evoked EPSCs occurring in the last 20 ms of the Masker stimulus. The two distributions differ significantly (p < 0.0001, Kolmogorov-Smirnov test). Average sEPSC amplitude of −97.28 ± 2.22 pA (median 82.10 pA) vs average eEPSC amplitude of −135.8 ± 3.24 pA (median 120.0 pA). Panels A-E show individual data points with the median and interquartile range overlaid (line). Synapses were classified as Δ pillar or ❍ modiolar, and as Low SR < 1 sEPSC/s ≤ High SR.

**Figure 2–figure supplement 1.**
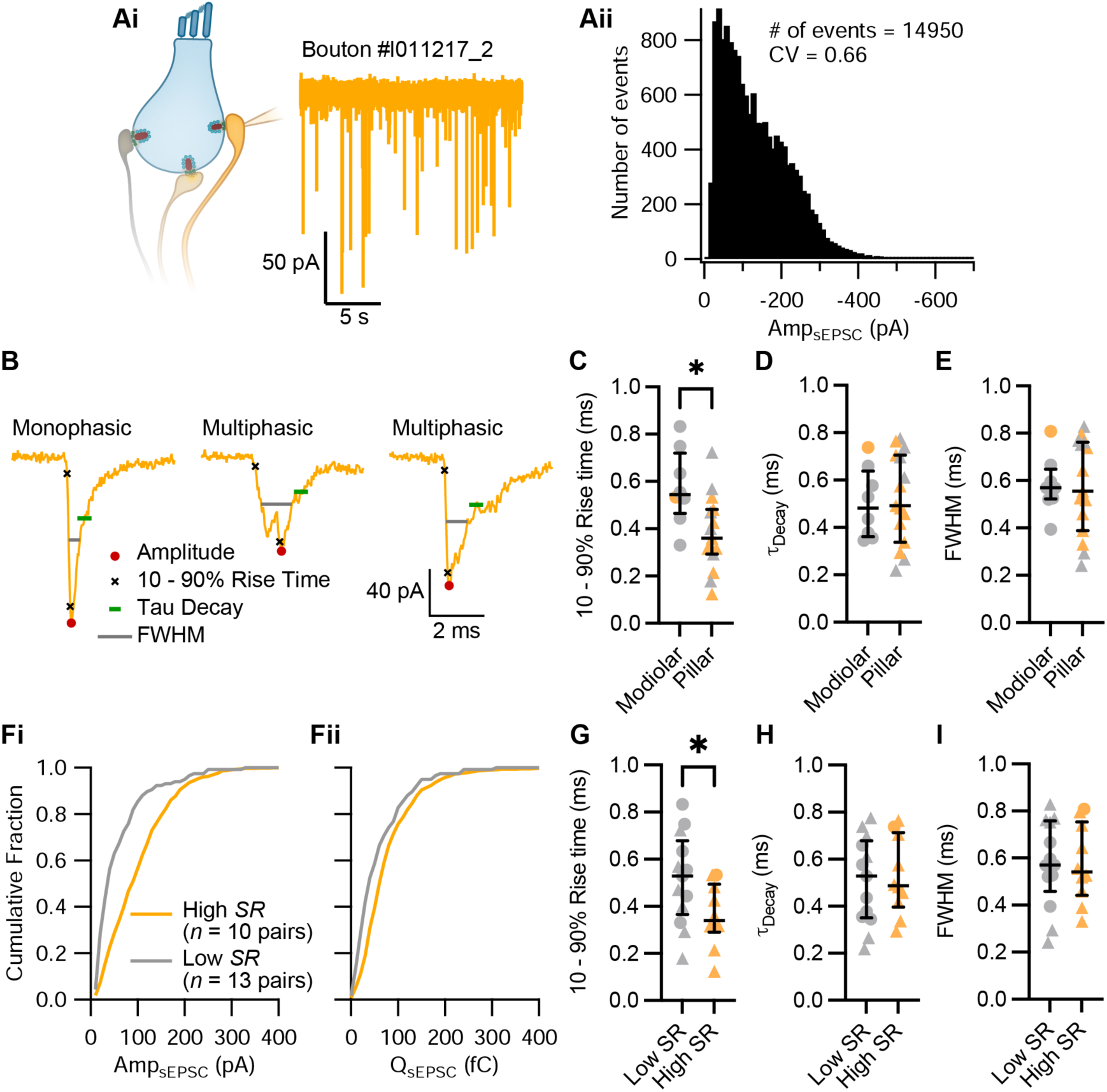
Pillar and high *SR* synapses have sEPSCs with faster rising times. **Ai-Aii.** Ruptured-patch whole-cell recordings from postsynaptic SGN boutons (without the IHC) displayed spontaneous rates from 0 up to 16.33 sEPSC/s. Pooled sEPSC amplitude (Aii) show a peak at −40 pA, with amplitudes ranging from around −10 pA to −670 pA. Bin size: 10 pA. **B.** Kinetics of sEPSCs, such as amplitude, 10-90% rise time, time constant of decay (τ_decay_) and full-width half-maximum (FWHM), were calculated with Neuromatic (Rothman and Silver, 2018). **C-E.** Pillar synapses had faster 10-90% rise times (C) than modiolar synapses, while their τ_decay_ (D) and FWHM (E) were comparable. **Fi-Fii.** Cumulative fraction of amplitude and charge of sEPSCs from low and high *SR* synapses. **G-I.** High *SR* synapses had faster 10-90% rise times (G) than modiolar synapses, while their τ_decay_ (H) and FWHM (I) were comparable. Panels C-E, G-I show individual data points with the median and interquartile range overlaid (line). Synapses were classified as △ pillar or ❍ modiolar, and as Low *SR* < 1 sEPSC/s ≤ High *SR*.

**Figure 3–figure supplement 1.**
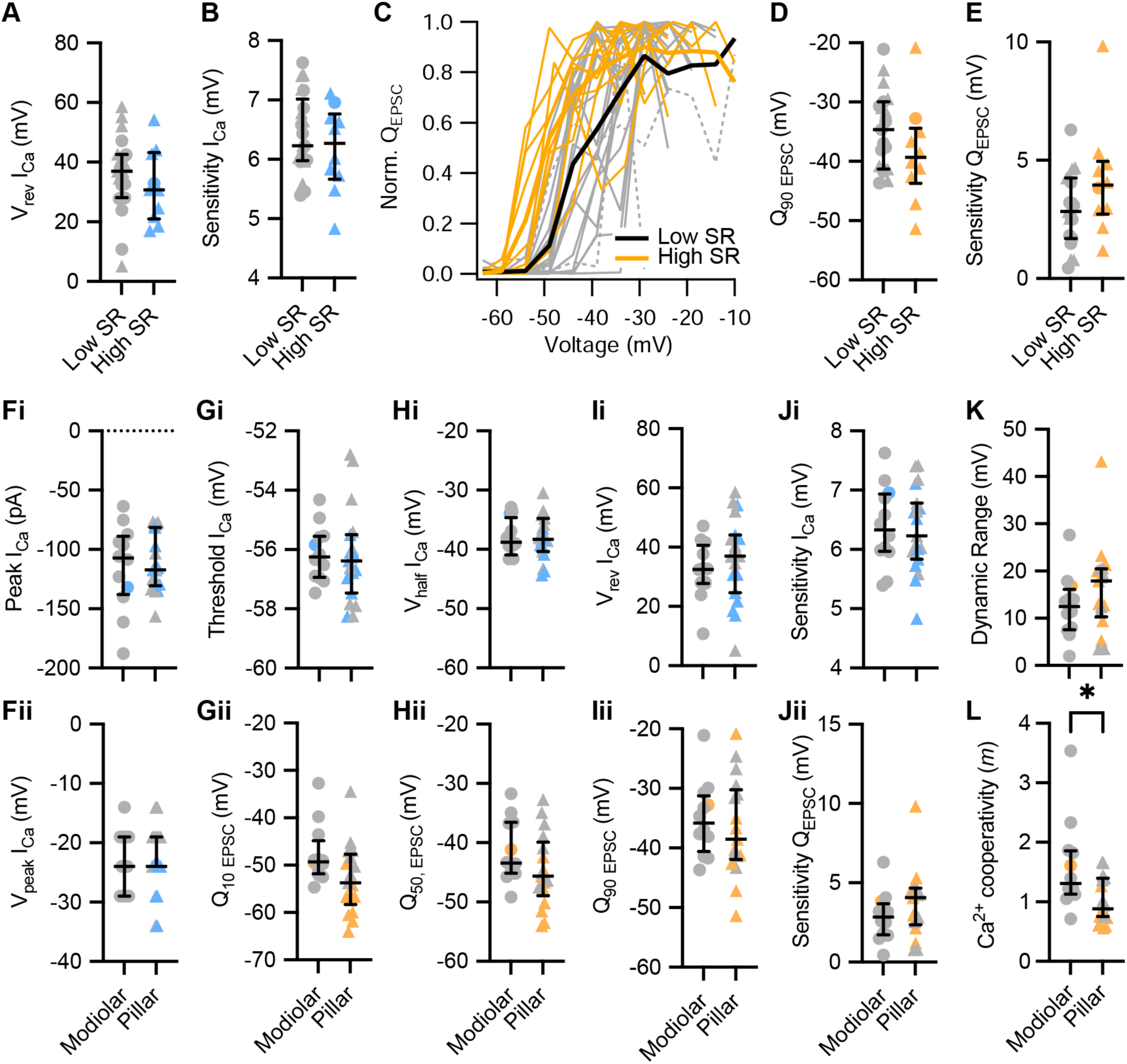
The voltage dependence of synaptic release does not differ significantly between modiolar and pillar synapses, but the Ca^2+^ dependence does. **A, B.** The reversal potential (A) and the voltage sensitivity of I_Ca_ (B) were comparable between low and high *SR* synapses. **C.** Triggered single active zone eEPSCs (release intensity curves) of 31 pairs of low and high *SR*. Averages (thick lines) and individual curves (thin lines) are overlaid. The release-intensity curve of two low *SR* pairs could not be fitted with the sigmoidal function (grey dotted lines). **D-E.** Q_90 EPSC_ (D) and voltage sensitivity of the release (E) were comparable between high and low *SR* synapses. **Fi-Fii.** The peak of Ca^2+^ current (Fi) and the voltage eliciting maximum Ca^2+^ current (Fii) were comparable between modiolar and pillar synapses. **Gi-Ji.** The threshold of Ca^2+^ influx (Gi), V_half_ I_Ca_ (Hi), reversal potential of I_Ca_ (Ii) and voltage sensitivity of I_Ca_ (Ji) did not differ between modiolar and pillar synapses. **Gii-Jii.** Q_10 EPSC_ (Gii), V_half_ Q_EPSC_ (Hii), Q_90 EPSC_ (Iii) and voltage sensitivity of the release (Jii) were comparable between modiolar and pillar synapses. **K.** Dynamic range of release was comparable between modiolar and pillar synapse. **L.** Ca^2+^ cooperativity (*m*) estimated for each individual synapse was significantly lower in pillar synapses. All panels but C show individual data points with the median and interquartile range overlaid (line). Synapses were classified as △ pillar or ❍ modiolar, and as Low *SR* < 1 sEPSC/s ≤ High *SR*.

**Figure 3–figure supplement 2.**
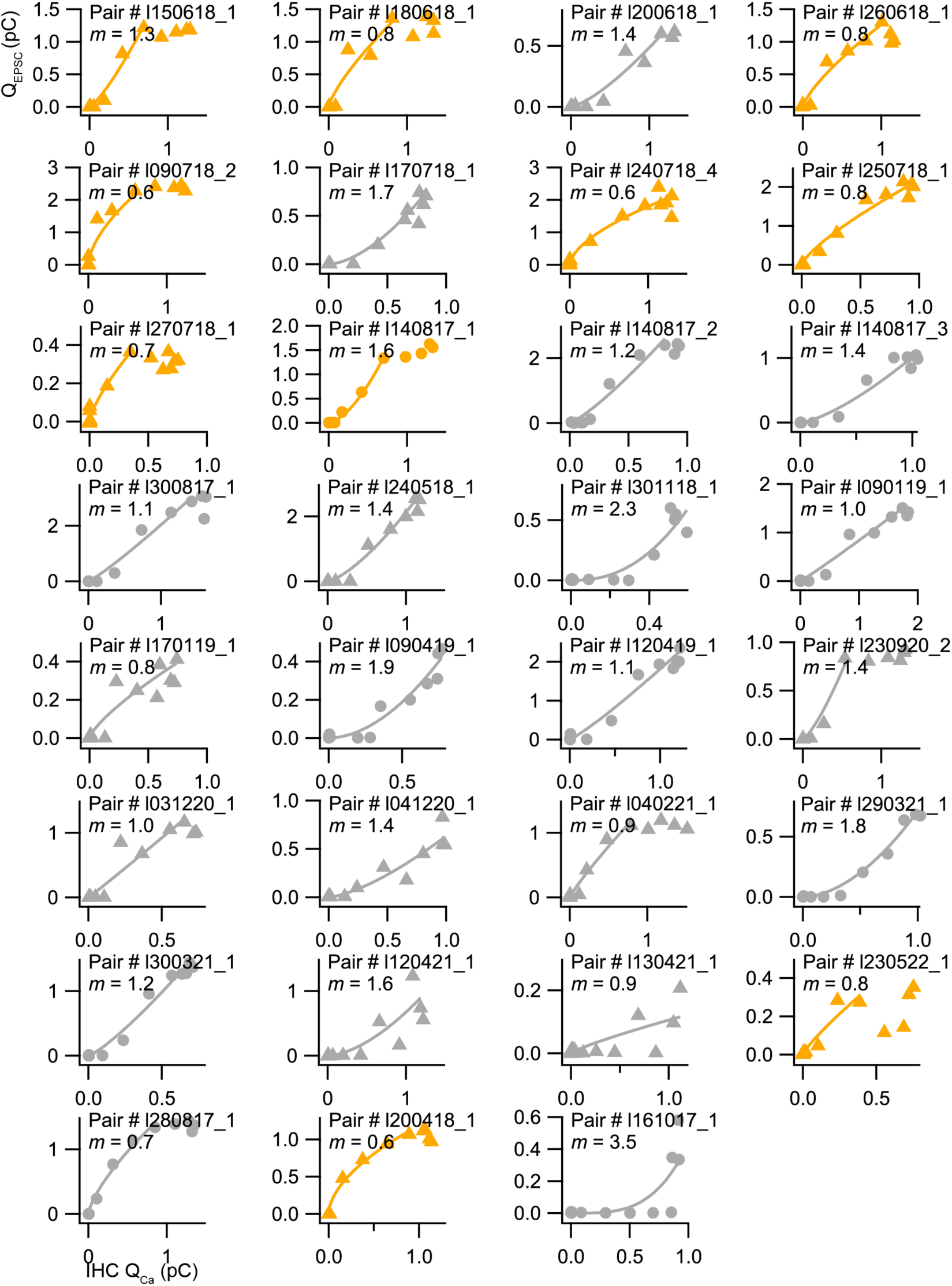
Apparent Ca^2+^ dependence of neurotransmitter release at individual synapses in the range of IHC receptor potentials. Scatter plots of the EPSC charges (Q_EPSC_) vs. the corresponding Ca^2+^ current integrals (Q_Ca_) for each individual synapse in response to 10 ms depolarizations from −58 to −19 mV. The solid line is a least-squares fit of a power function (Q_EPSC_ = a(Q_Ca_)*^m^*) to each pair data. Synapses were classified as △ pillar or ❍ modiolar, and as Low *SR* < 1 sEPSC/s ≤ High *SR*.

**Figure 4–figure supplement 1.**
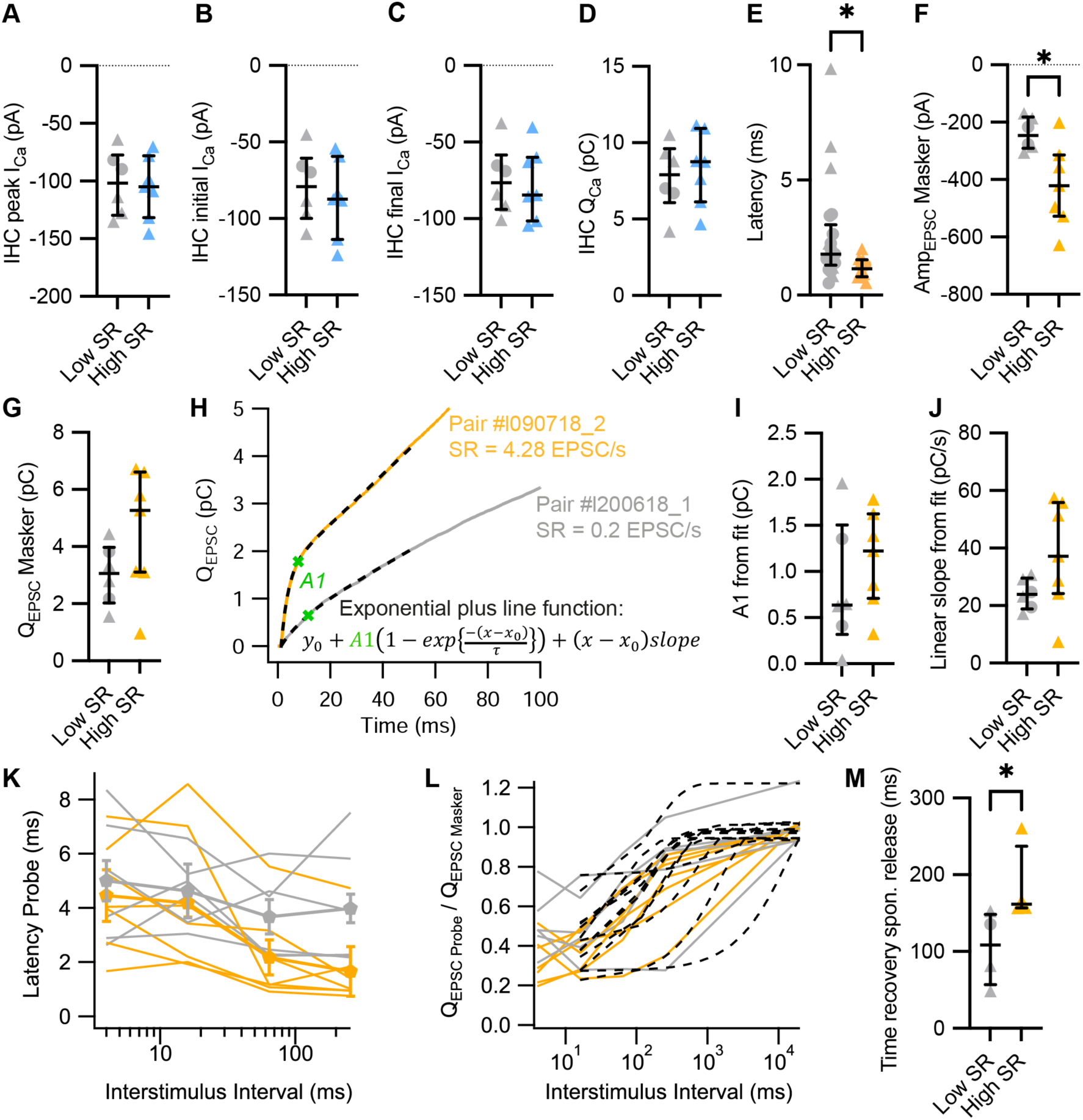
Parameters of synaptic vesicle pool dynamics in forward masking protocols. **A-D.** Presynaptic IHC peak (A), initial (B) and final (C) Ca^2+^ current (I_Ca_), and Ca^2+^ charge (Q_Ca_) during the masker stimuli were comparable regardless the *SR* of postsynaptic bouton. **E.** Latencies of the evoked EPSCs were also significantly shorter in high *SR* than low *SR* synapses when compared in a bigger sample size (31 pairs from Fig. 4). **F-G.** High *SR* synapses had significantly larger amplitudes of evoked EPSCs in response to the masker stimulus (F), while the EPSC charge (Q_EPSC_) did not differ (G). **H.** Exemplary average Q_EPSC_ response of a high *SR* and a low *SR* synapse to the masker stimuli. We fitted an exponential plus line function to the first 50 ms of the response (discontinuous lines) to study SV pool depletion dynamics. From these fits, we can retrieve information about RRP size (amplitude of the exponential component, *A1*), RRP depletion time constant (τ of the exponential component) and sustained release (linear slope of the line component). **I-J.** Amplitude (I) and linear slope (J) obtained from the fits of the exponential plus line function to the Q_EPSC_ from individual pairs (Fig. 4E). **K.** Latencies of the response to the probe stimuli (EPSC_onset_ - Probe_onset_) for different ISI. **L.** Single exponential fits from 16 ms to 20000 ms (black dotted lines) to estimate the recovery kinetics from RRP depletion. **M.** The spontaneous release after the probe offset recovered slower in high *SR* synapses. Panels A-G, I, J, M show individual data points with the median and interquartile range overlaid (line). Synapses were classified as △ pillar or ❍ modiolar, and as Low *SR* < 1 sEPSC/s ≤ High *SR*.

**Supplementary Table 1.** Spontaneous activity (*SR*) was calculated from time windows without stimulation with the IHC held at −58 mV (Total time for SR calculation). This total time was calculated from the cumulative recording time of either from 5 – 10 s recordings and/or from the segments before and after a depolarizing pulse.

| Cellname | Side | SR | Total time for SR calculation | Cumulative time of 5-10 s recordings (s) | Cumulative time before/after a depolarizing pulse (s) |
| --- | --- | --- | --- | --- | --- |
| I140817_1 | Modiolar | 1.07 | 36 |  | 36 |
| I140817_2 | Modiolar | 0.38 | 28.8 |  | 28.8 |
| I140817_3 | Modiolar | 0.05 | 43.2 |  | 43.2 |
| I300817_1 | Modiolar | 0.00 | 31.2 |  | 31.2 |
| I240518_1 | Pillar | 0.22 | 91.2 |  | 91.2 |
| I180618_1 | Pillar | 1.00 | 126.8 |  | 126.8 |
| I031220_1 | Pillar | 0.00 | 10 | 10 |  |
| I041220_1 | Pillar | 0.00 | 10 | 10 |  |
| I280817_1 | Modiolar | 0.00 | 3.69 |  | 3.69 |
| I200418_1 | Pillar | 3.92 | 3.06 |  | 3.06 |
| I200618_1 | Pillar | 0.20 | 20.8 |  | 20.8 |
| I260618_1 | Pillar | 2.05 | 20.8 |  | 20.8 |
| I090718_2 | Pillar | 4.28 | 56.8 |  | 56.8 |
| I170718_1 | Pillar | 0.38 | 23.2 |  | 23.2 |
| I240718_4 | Pillar | 17.93 | 7.2 |  | 7.2 |
| I250718_1 | Pillar | 3.18 | 38.4 |  | 38.4 |
| I270718_1 | Pillar | 9.00 | 20 |  | 20 |
| I090119_1 | Modiolar | 0.00 | 20 | 20 |  |
| I150618_1 | Pillar | 1.49 | 12 |  | 12 |
| I290321_1 | Modiolar | 0.03 | 38 | 10 | 28 |
| I300321_1 | Modiolar | 0.36 | 85.2 | 50 | 35.2 |
| I120421_1 | Pillar | 0.21 | 35.6 | 10 | 25.6 |
| I130421_1 | Pillar | 0.00 | 35.4 | 25 | 10.4 |
| I301118_1 | Modiolar | 0.15 | 40 | 40 |  |
| I170119_1 | Pillar | 0.80 | 10 | 10 |  |
| I090419_1 | Modiolar | 0.00 | 10 | 10 |  |
| I120419_1 | Modiolar | 0.70 | 10 | 10 |  |
| I230920_2 | Pillar | 0.00 | 15 | 15 |  |
| I040221_1 | Pillar | 0.20 | 10 | 10 |  |
| I230522_1 | Pillar | 1.26 | 49.6 | 10 | 39.6 |
| I230817_2 | Modiolar | 0.16 | 6.12 |  | 6.12 |
| I161017_1 | Modiolar | 0.00 | 2.34 |  | 2.34 |
| I200522_2 | Pillar | 0.00 | 40 | 40 |  |

